# Distinct Functional States of Neutrophils by Actin Disassembly and NF-kB/STAT3 Signaling

**DOI:** 10.1101/2023.07.14.548975

**Authors:** Niko Heineken, Jan C. Schumacher, Tarik Exner, Friederike S. Neuenfeldt, Jüri Habicht, Franz Kahlich, Hadrian Platzer, Uta Merle, Tobias Renkawitz, Yvonne Samstag, Ricardo Grieshaber-Bouyer, Stella Autenrieth, Guido Wabnitz

## Abstract

Polymorphonuclear neutrophils (PMNs) can differentiate into distinct states, which can either exacerbate or resolve inflammation. Our study shows that mice challenged with TLR agonists exhibited PMN differentiation along two major paths characterized by the expression of CCR5 or PD-L1. Similar differentiation was observed in the blood of severe COVID-19 patients and the synovial fluid of osteoarthritis patients. Prolonged *in vitro* priming of human PMNs modeled the differentiation paths. Actin disassembly favored CCR5 upregulation, while NF-kB activation stabilized the actin cytoskeleton and suppressed the development of CCR5^+^ PMNs. Additionally, PD-L1 upregulation was triggered by STAT3 signaling and NF-kB activation. Functionally, CCR5 expressing PMNs were pro-NETotic, while PD-L1^+^ PMNs showed immunosuppressive functions by inhibiting T cell proliferation via PD1. Together, PMN differentiation depended on the priming conditions, and the balance between actin disassembly and NF-kB/STAT3 activation translated the present micro-milieu into phenotypic and functional diversification of PMNs.

**Synopsis:** Neutrophils underwent phenotypical and functional diversification both *in vivo* and *in vitro*. Actin disassembly led to the generation of CCR5^high^ neutrophils with increased spontaneous NETosis, whereas NF-kB and STAT3 induced PD-L1 expression with T-cell suppressive properties as a deviation from the default pathway.

- PMN of mice challenged with TLR agonists develop two distinct phenotypes, CCR5^high^ and PD-L1^high^.
- CCR5 and PD-L1-defined neutrophil phenotypes were found in blood of patients with severe COVID-19 and in the synovial fluid of osteoarthritis patients.
- *In vitro* priming induced a similar bifurcation of PMN phenotypes marked by either CCR5 or PD-L1.
- Actin disassembly preceded canonical development of CCR5^+^ PMN.
- NF-kB halted actin disassembly by LPL regulation.
- During neutrophil priming, STAT3 aided NF-kB in the expression of PD-L1.

**Graphical abstract:** 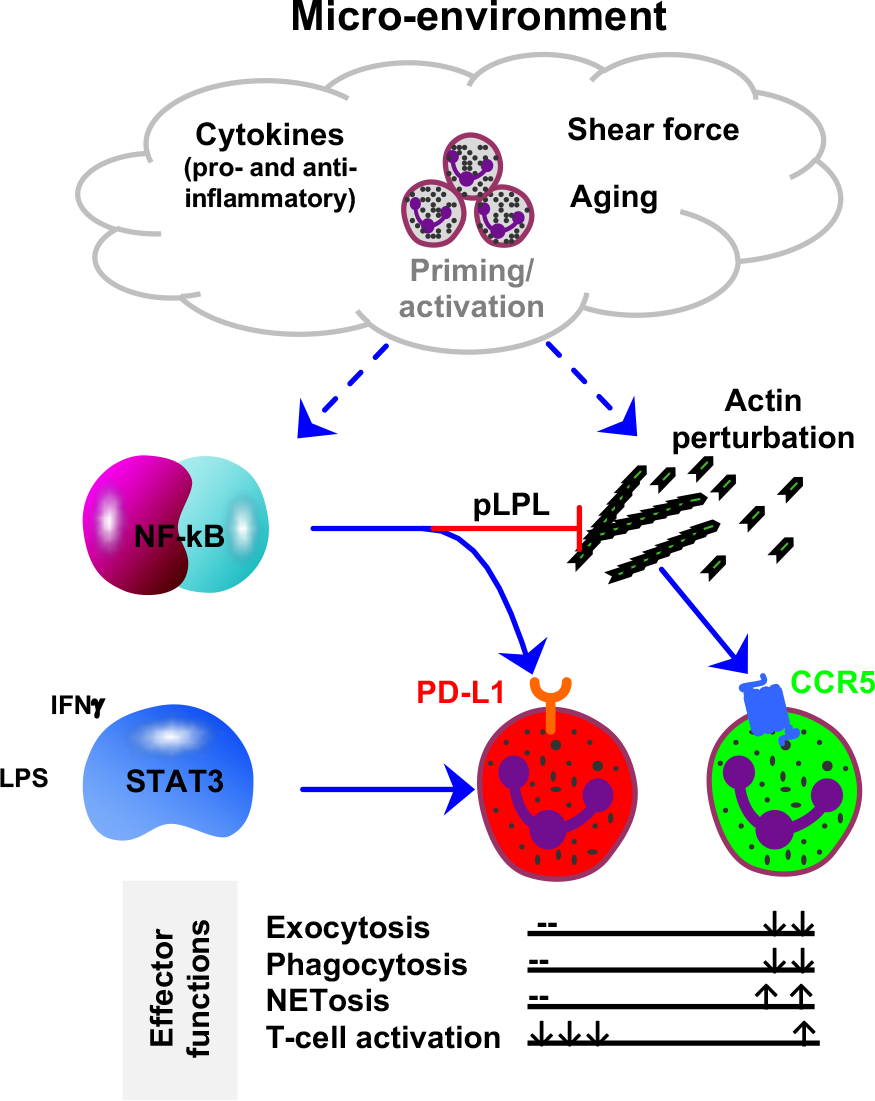

## Introduction

Polymorphonuclear neutrophils (PMNs) play a crucial role in immunity and inflammation. PMN priming, e.g. by bacterial products or cytokines like GM-CSF, TNF or INFγ, is a critical factor in inflammation and infections as it renders the cells more responsive to pathogens and lowers their activation threshold (Condliffe et al., 1998; Ekpenyong et al., 2017; Elbim et al., 1994; Hallett and Lloyds, 1995; Liu et al., 2017; Miralda et al., 2017; Potera et al., 2016). Moreover, it has been shown that PMN subgroups or polarization states exist in inflammation. Several approaches have been used for the analysis of PMN phenotypes *in vivo* including multi-omics analysis of PMNs from different tissues under physiological conditions, infection, or inflammation (Ballesteros et al., 2020; Grieshaber-Bouyer et al., 2021; Xie et al., 2020). The mechanisms underlying diversification of mature PMN in response to the microenvironment are not well understood (Deniset and Kubes, 2018; Grieshaber-Bouyer et al., 2021; Neuenfeldt et al., 2022b; Pillay et al., 2012; Soehnlein et al., 2017; Xie et al., 2020).

So far, the priming of PMNs has been experimentally mimicked *in vitro* by short-term incubations with low concentrations of the priming agents (Lamb et al., 2012; Wright et al., 2013; Yan et al., 2019). However, *in vivo*, PMNs can be exposed to the prevailing micro-milieu for prolonged periods, and cytokine compositions and concentrations can vary widely. Therefore, in this study, we compared PMN phenotypes occurring *in vivo* in mice and human to those generated *in vitro*. We examined the diversification of PMNs from the spleens of mice treated with TLR agonists and compared these with PMN subsets present in patients either infected with SARS-CoV2 and exhibiting flourishing systematic inflammation or in the joints of patients with osteoarthritis, associated with chronic low-grade inflammation. Experiments with isolated human PMNs were performed *in vitro* to simulate inflammatory settings in humans. For *in vitro* priming we selected bacterial products (LPS and fMLP) and cytokines based on their known role in inflammatory processes. We included combinations of different cytokines, since these are particularly effective for PMN priming and better mimic the condition *in vivo* (Mol et al., 2021). Dimensionality reduction and tree-shaped trajectories were applied to model PMN heterogeneity, and functional properties of the resulting PMN subpopulations were investigated.

In all the approaches utilized, significant phenotypic diversification of PMNs was observed. Of particular interest was the emergence of two distinct subgroups distinguished by the expression of CCR5 (CD195) or PD-L1 (CD274). CCR5^+^ PMNs with enhanced spontaneous NETosis and low-level T cell activation capabilities developed as canonical pathway and were still present after TNF priming, while IFNγ or LPS skewed PMN diversification towards PD-L1^+^ PMNs with strong phagocytosis capability and T cell suppressive functions. The equilibrium between actin disassembly and NF-kB/STAT3 activation played a pivotal role in determining the phenotype of PMNs. Together, this study offers significant insights into the complex process of PMN diversification under different microenvironmental conditions. By examining the dynamic nature of PMN phenotypes, our findings contribute to the understanding of the intricate interplay between PMN subpopulations and cytokines, deepening our knowledge of the underlying mechanisms involved in inflammatory responses.

## Results

### PMN diversification after challenging mice with TLR agonists

Age and weight-matched wild-type mice were treated with tail vein injections of sublethal doses of *S. typhimurium* LPS (TLR4 priming), the synthetic triacylated lipopeptide Pam3CSK4 (TLR2 priming) or the viral analog Poly(I:C) (TLR3 priming). PMNs were isolated from the spleen after 24h and analyzed by spectral flow cytometry (**Fig. 1A, upper part**). PMNs were gated as CD11b^+^Ly6G^+^ and Ly6C^int^ events (**Fig. 1A, lower part**). Gated events and their median fluorescence intensity values were concatenated and classified into 36 different clusters using the SOM (self-organizing maps) algorithm (**Fig. S1A**). To further summarize the data included in the 36 clusters unbiased branching analysis combined with hierarchical cluster analysis (HCA) was performed revealing 7 meta-clusters (**Fig. 1B**). To better visualize the distribution of meta-clusters between conditions, we created UMAP representations showing that the distribution of PMNs on the UMAP landscape differed strongly after LPS, Pam3CSK4, or Poly(I:C) challenge (**Fig. 1C**). Meta-cluster 3, 5, 6 and 7 were the main meta-clusters in control (PBS) mice. LPS conditions induced a shift toward meta-cluster 2, and Poly(I:C) and Pam3CSK4 towards 1 and 4. Considering the expression patterns of individual meta-clusters, it was evident that CCR5 (CD195) showed a stronger signal only in meta-cluster 2. Meta-cluster 1 and 4 were characterized by expression of PD-L1 (CD274). Applying MST (minimal spanning tree) projection created with this dataset and colored by assigned meta-clusters revealed a relationship between meta-clusters 1 and 4, which were separated from meta-cluster 2 verifying CCR5 and PD-L1 as markers for distinguishing PMN phenotypes (**Fig. 1D and S1B**). CCR5^+^ and PD-L1^+^ and double positive (DP) PMNs could be detected by downstream analysis applying traditional biaxial gating (**Fig. 1E**). After the administration of LPS, there was a notable rise in the abundance of CCR5^+^ PMNs. However, treatment with Poly(I:C) or Pam3CSK4 resulted in a substantially greater quantity of PD-L1^+^ PMNs (**Fig. 1F-G**). This implies that the ratio of specific PMN phenotypes is context-dependent.

**Figure 1.**
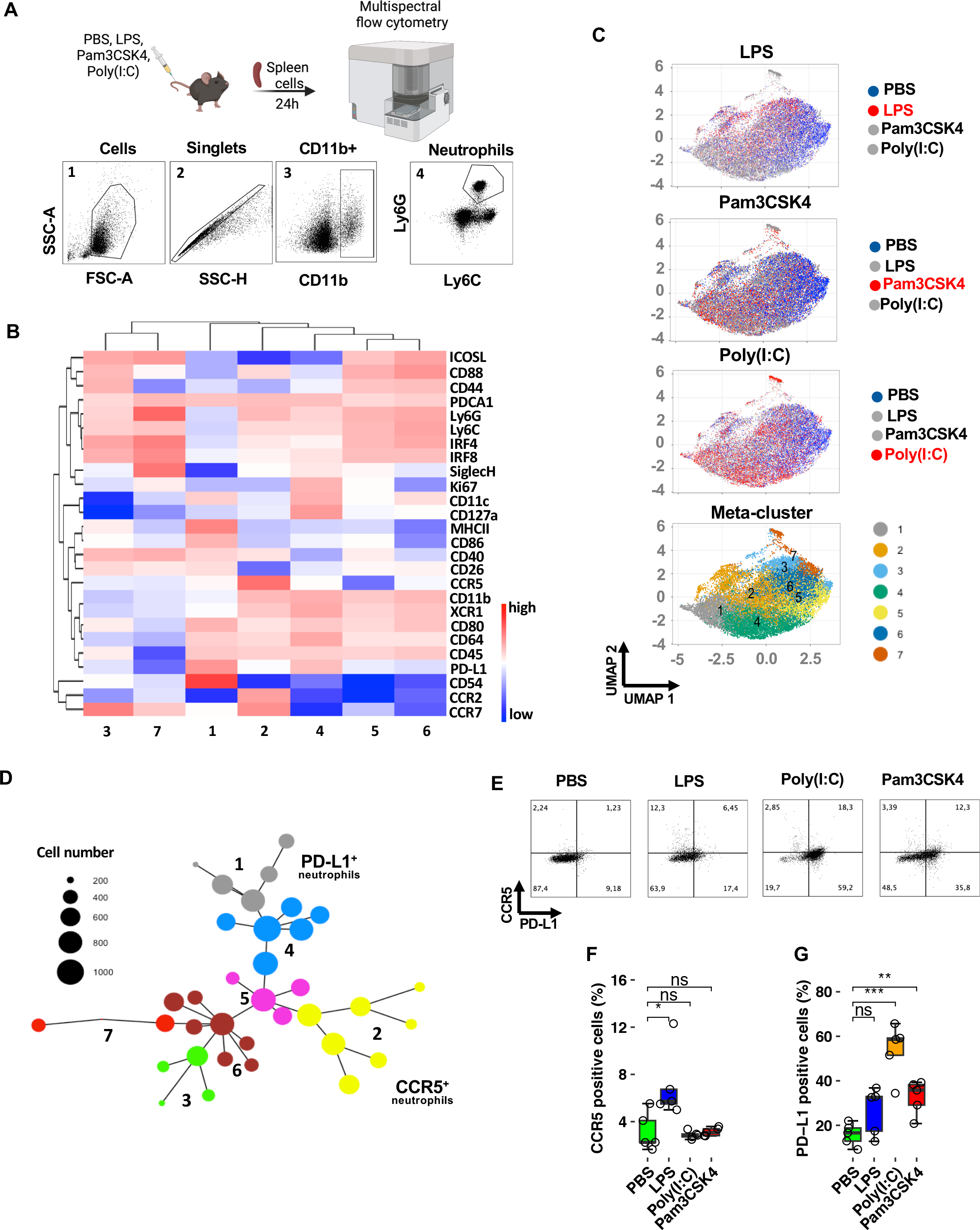
Divers phenotypes of PMNs in challenged mice. **A**) Phenotypic characterization of PMNs in challenged mice. Mice were treated with PBS, LPS, Pam3CSK4, or Poly(I:C) for 24 h. Splenocytes were stained with a 33-color antibody panel and analyzed by spectral flow cytometry. PMNs were identified as CD45^+^CD11b^+^Ly6G^+^Ly6C^int^. **B**) Hierarchical branch heatmap generated using CytoTree analysis. The heatmap represents the clustering of PMN phenotypes from 5 mice for each condition. **C**) UMAP projection analysis using CytoTree, showing merged samples of PMNs from mice injected with PBS, LPS, Pam3CSK4, or Poly(I:C) into the tail vein (n=5 per condition). The plots depict single-cell flow cytometry-based receptor expression data, with color-coded branches (lower part) or conditions (upper part). **D**) Construction of PMN diversification trajectory based on UMAP coordinates using minimum spanning tree (MST) analysis. Branch IDs were annotated manually. **E**) Biaxial gating showing the percentage of CCR5^+^ and PD-L1^+^ PMNs across conditions as indicated (representative of 5 mice per condition). **F-G**) Quantification of CCR5^+^ and PD-L1^+^ PMNs, as manually gated (n=5, SEM, ANOVA, n.s. = not significant, * p<0.05, ** p<0.01, *** p<0.001). The original gating strategy is presented in supplementary Figure 1C.

### Diversified PMNs are evident in COVID-19 and osteoarthritis patients

TLRs are important during SARS-CoV-2 infections (Sariol and Perlman, 2021). Coronavirus disease 2019 (COVID-19) can range from asymptomatic to severe with acute respiratory distress syndrome (ARDS). The latter is associated with a cytokine storm, in particular high levels of interferons (Lee and Shin, 2020; Mehta et al., 2020; Todorovic-Rakovic et al., 2022). Therefore, we examined PMNs from the blood of patients with severe COVID-19 infections and convalesce patients (3 months after discharge from the intensive care unit). We observed a significant upregulation of CD66b on PMNs of COVID-19 patients, confirming their activation (**Fig. 2A and B**). Although the expression of CD66b on PMNs from recovered individuals had dropped, a significant increase in expression was still detected compared to PMNs from healthy donors. Both PD-L1^+^ and DP PMNs were expanded in the blood of the COVID-19 patients (**Fig. 2C-D**). Clustering analysis and heatmapping also identified populations that showed dominant expression of PD-L1 and CCR5 (meta-cluster 2 and 3) (**Fig. 2E**). Both populations were double positive. Meta-cluster 5 and 1 also expressed higher amounts of PD-L1 compared to meta-cluster 4. The presence of the latter was primarily observed in the control and convalesce group, as indicated by MST analysis (**Fig. 2F**).

**Figure 2.**
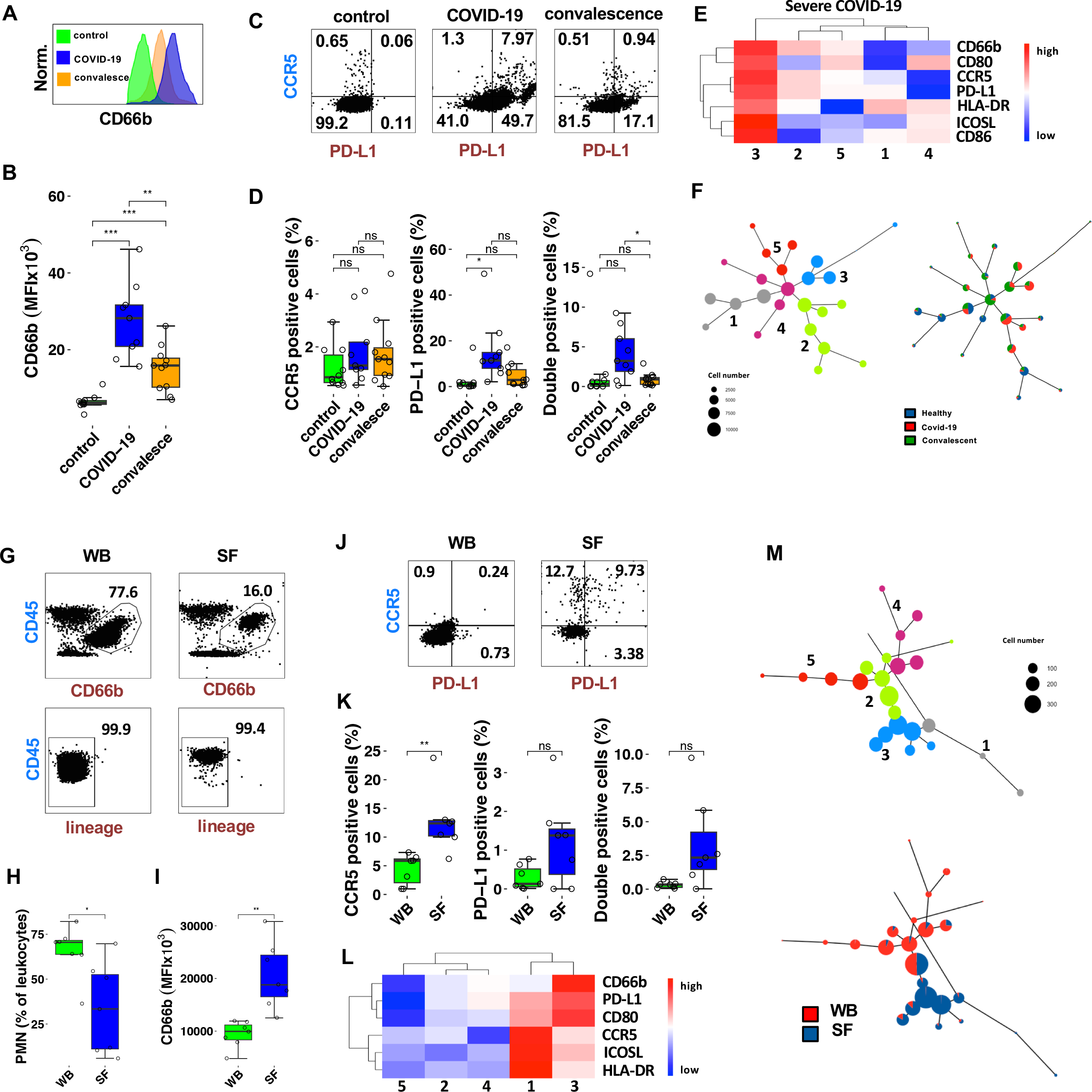
PMN phenotypes in COVID-19 and osteoarthritis patients. **A-B**) CD66 expression on PMNs in different cohorts (n=10 per condition): COVID-19 patients during their ICU stay, convalescent COVID-19 patients (3 months post-recovery), and healthy control individuals. PMNs were stained in whole blood and analyzed by flow cytometry. (A) Representative histogram depicting CD66 expression. (B) Quantification of median fluorescence intensities (MFIs) from 10 individuals per group. **C-D**) Expression of CCR5 and PD-L1 was determined by flow cytometry on peripheral blood PMNs from the same cohorts as described in A. While C shows representative examples, D contains a statistical evaluation (n=10 per condition, SEM, ANOVA, n.s. = not significant, * p<0.05). **E-F**) Hierarchical branch map (E) and tree plots (MSTs) showing branch positioning (F, left part) or conditions (right part). **G**) Gating strategy to identify PMNs from whole blood (WB) or diluted synovial fluid (SF) of osteoarthritis patients. **H-I**) Quantification of relative PMN amount measured as percent of leukocytes (H), and expression of CD66b on PMNs (I) (n=6, SEM, Mann Whitney U test, * p<0.05). **J-K**) Flow cytometry was performed to analyze the expression levels of CCR5 and PD-L1 on PMNs. Representative flow cytometry plots are shown in J. Statistical evaluation was conducted on 6 patients. Error bars represent the standard error of the mean (SEM). Statistical significance was determined using Mann Whitney U test (n.s. = not significant, * p<0.05). **L**) Hierarchical branch heatmap generated using CytoTree analysis representing PMNs derived from peripheral blood and synovial fluid of the described osteoarthritis (OA) patients. The heatmap illustrates the clustering of PMN phenotypes based on flow cytometry data. **M**) Constructions of the PMN diversification trajectory using minimum spanning tree (MST) analysis are shown. The trajectories are colorized based on branches (upper part), indicating distinct developmental paths of PMN subsets, and PMN origin (lower part), distinguishing between peripheral blood and synovial fluid PMNs.

We next analyzed PMN of osteoarthritis (OA) patients. OA and COVID-19 differ significantly in terms of inflammation and localization, representing two distinct diseases. This enables us to make conditional statements regarding the occurrence of PMN phenotypes under varying conditions. Furthermore, it allows the comparison of matched PMNs obtained from the synovial fluid (SF) and whole peripheral blood (WB) of the same patient. These fluids were collected during elective surgery from OA patients who had joint prostheses implanted. PMNs were identified in these samples by flow cytometry using the SSC profile, expression of CD45 plus CD66b, and absence of the lineage markers (**Fig. 2G**). The percentage of PMNs out of all leukocytes was significantly smaller in synovial fluid than in the peripheral blood of the matched OA patients (**Fig. 2H**). However, greatly increased CD66b expression on SF-PMNs indicated activation of the cells (**Fig. 2I**). While PMNs were homogeneous in peripheral blood, SF-PMNs showed diversification into CCR5^+^, PD-L1^+^, and double-positive PMNs (**Fig. 2J**). Across all patients, CCR5^+^ PMNs in particular showed a significant increase in their frequency (**Fig. 2K**). A branch heatmap verified that both CCR5^+^ PMNs (meta-cluster 5) and CCR5^+^/PD-L1^+^ double-positive PMNs (meta-clusters 4 and 6) were present in osteoarthritis synovial fluid (**Fig. 2L**). Representation as a tree revealed that CCR5^+^ PMNs (meta-cluster 5) and double-positive PMNs (meta-clusters 4 and 5) were mainly present in SF (**Fig. 2M**). Together, these results demonstrate that CCR5^+^ and PD-L1^+^ neutrophils arise in distinct inflammatory contexts *in vivo* and that the proportion of these cells differs depending on the specific inflammatory setting.

### PMN diversification paths of *in vitro* primed human PMNs

To understand which pathways are associated with the diversification of PMNs, we tested whether diversification of primary human PMNs can also be induced *in vitro*. While significant cell death can occur depending on cell culture conditions, cytokines significantly extend the short life span of PMNs (Cui et al., 2021; Derouet et al., 2004; Kolaczkowska and Kubes, 2013). With GM-CSF plus IFNγ priming, only less than 5% of the cells were Zombie positive after 24h (**Fig. S2A**). In addition, we measured nuclei fragmentation using imaging flow cytometry (**Fig. S2B**) (Balta et al., 2017; Lowman et al., 2010; Neuenfeldt et al., 2022a; Wabnitz et al., 2010a) and membrane flip with Apotracker-binding verifying the results achieved with Zombie-staining (**Fig. S2C**).

To analyze phenotypic alterations, freshly isolated PMNs from the peripheral blood of healthy individuals were analyzed immediately after purification (resting-state PMNs, rPMNs) or after incubation with the bacterial products LPS or fMLP as well as after priming with different pro-inflammatory cytokine regimes, i.e. GM-CSF, IFNγ, IL-8, a combination of IFNγ and GM-CSF as well as TNF or the anti-inflammatory cytokine IL-10 for 24 h. Note that GM-CSF, IFNγ, IL-10 and TNF were also present in the synovial fluid of osteoarthritis patients. (**Fig. S3**). Cells were stained with a 7-color flow-cytometry panel including anti-CCR5 and anti-PD-L1 antibodies. Higher numbers of Zombie^+^ cells were only detected in control conditions, i.e. in the absence of priming agents (>20% Zombie^+^ cells), or after priming with fMLP, IFNγ, IL-8 and IL-10 (>10% Zombie^+^ cells) (**Fig. S2A**). A significant amount of Zombie^-^ PMNs expressed CCR5 under control conditions, while PD-L1 upregulation was not induced (**Fig. 3A and B**). Priming with various cytokines or incubation with bacterial products led to a decrease in the CCR5^+^ PMN subpopulation compared to the control conditions. PD-L1^+^ PMNs were found in conditions that included IFNγ or LPS and were highly abundant when cytokine combinations (GM-CSF + IFNγ) were applied. Here, also CCR5/PD-L1 double-positive cells were present.

**Figure 3.**
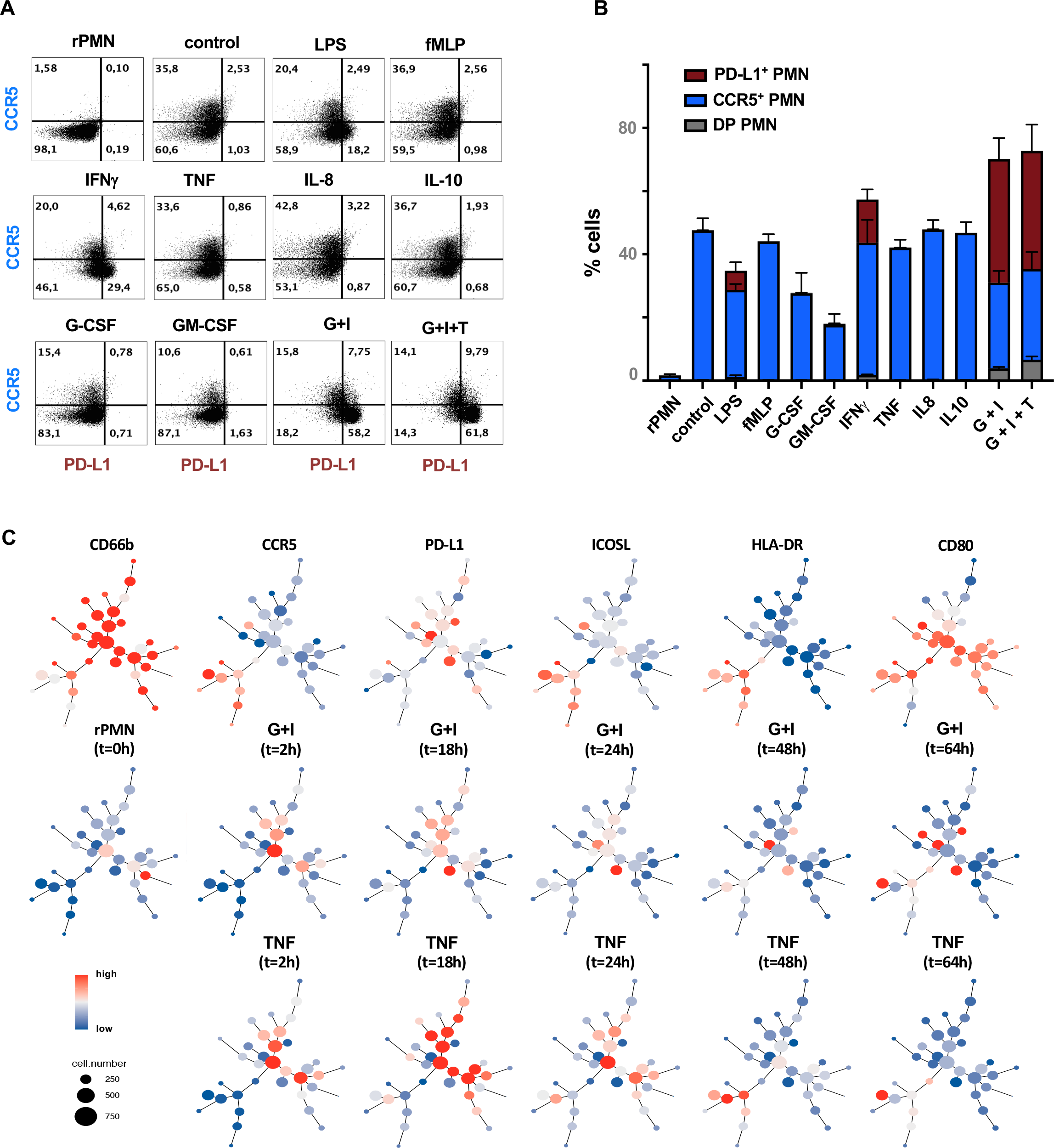
*In vitro*-induced PMN diversification paths. **A-B**) Flow cytometric analysis of CCR5 and PD-L1 expression on live PMNs primed *in vitro* using the indicated compounds (A). The time point is 24h. The compound combination G+I represents GM-CSF + IFNγ, while G+I+T represents GM-CSF + IFNγ plus TNF. Quantification of single positive and double positive (DP) cells, including standard error of the mean (SEM), is shown in B. **C**) Construction of PMN diversification trajectory based on UMAP coordinates using MST calculated on a time course analysis as indicated. The upper trees show the receptor distribution as expression heatmap. The middle (G+I) and lower (TNF) tree rows show temporal superimpositions according to the indicated time points and priming conditions. In the lower two rows, cell numbers corresponding to the specific condition are represented through color coding.

We next focused on distinguishing between classification and established subgroups of PMNs characterized by expression of CD62L, CD16 and CD177 (Silvestre-Roig et al., 2016). Using an extended flow-cytometry panel and GM-CSF plus IFNγ or TNF priming showed that CCR5^+^ PMN were mainly CD62L^low^/CD16^low^ whereas PD-L1^+^ PMN contained both CD62L^high^/CD16^high^ and CD62L^low^/CD16^low^ (**Fig. S2D-G**). Considering the existence of two PMN subgroups in peripheral blood that exhibit different levels of CD177 expression (Eulenberg-Gustavus et al., 2017), we investigated if this dichotomy could be responsible for the presence of CCR5^+^ and PD-L1^+^ PMN. However, the development of CCR5^+^ and PD-L1^+^ PMN was independent on CD177 expression as both FACS-sorted CD177^+^ and CD177^-^ PMN gave rise to similar proportions of CCR5^+^ and PD-L1^+^ PMN upon GM-CSF plus IFNγ priming (**Fig. S2H**). In the subsequent set of experiments, time-course analysis was performed. PMNs were incubated with GM-CSF plus IFNγ or TNF and phenotypically characterized at different time points, i.e. 0, 2, 18, 24, 48 and 64 h (4 distinct biological samples per time point) to explore temporal diversification of PMNs. Note that the number of Zombie^+^ cells was low even after 48h but increased thereafter (**Fig. S2I**). After excluding Zombie^+^ cells and pre-processing all 41 samples, cells were classified into 36 different clusters (**Fig. S4A**). Furthermore, MSTs were computed to illustrate the clustering distribution. Cluster 31 defined the root cluster from which CCR5^+^ and PD-L1^+^ PMNs developed (**Fig. 3C and S4B**). It is evident that the CCR5^+^ PMNs were predominantly concentrated within a single branch and separated from PD-L1 expressing PMN (**Fig. 2C, upper part**). When observing temporal plots, it becomes apparent that cells diverge more rapidly from the root clusters following GM-CSF plus IFNγ treatment compared to TNF priming (**Fig. 2C, middle and lower part**). In addition, trajectory inference was utilized to reconstruct hierarchical transitions of PMNs and to establish the temporal ordering of individual cells based on their expression profiles, also known as “pseudotime” in computational inference (**Fig. S5**). By analyzing the correlation between pseudotime and marker expression, we also observed that PMNs acquired CCR5, HLA-DR and ICOSL progressively within the pseudotime trajectory, while CD66b and CD80 decreased. PD-L1 increased initially and then decreased slightly with progression through pseudotime.

### NF-kB and STAT3 regulate PMN diversification

To get more information about diversification paths, gene set analysis (GSA) of bulk mRNA expression data of rPMN or primed PMNs (GM-CSF plus IFNγ or TNF) were analyzed. Principal component analysis (PCA) revealed that gene expression profiles differ between GM-CSF plus IFNγ- or TNF-primed PMNs, rPMNs, or control cells (**Fig. S6A**). GSA z-scores as well as adjusted significances were calculated and projected on a radar plot (**Fig. 4A and supplement material 1**). The gene set “Cytoplasm” was most significantly enhanced in GM-CSF plus IFNγ-primed cells, whereas the “JAK-STAT cascade” gene set was equally expressed in rPMNs and GM-CSF plus IFNγ-primed cells and the “signal transduction” gene set was significantly increased in TNF-primed cells. PCA analysis of the gene set “Cytoplasm” verified distinctiveness across conditions (**Fig. S6B**). A detailed analysis of the “Cytoplasm” gene set showed that both types of priming (GM-CSF plus IFNγ and TNF) induced genes belonging to the NF-kB pathway (i.e. *RelA, NFkB1, NFKBI1, CHUK*) (**Fig. 4B**). In GM-CSF plus IFNγ-primed cells, genes from the JAK-STAT pathway (i.e. *STAT1* and *STAT3*) were also more strongly expressed than in TNF-primed PMNs. KEGG (Kyoto Encyclopedia of Genes and Genomes) analysis of gene expression data confirmed enhanced expression of the canonical NF-kB pathway in GM-CSF plus IFNγ- and TNF-primed PMNs and upregulation of JAK-STAT signaling in GM-CSF plus IFN-primed-primed cells (**Fig. S6C**).

**Figure 4.**
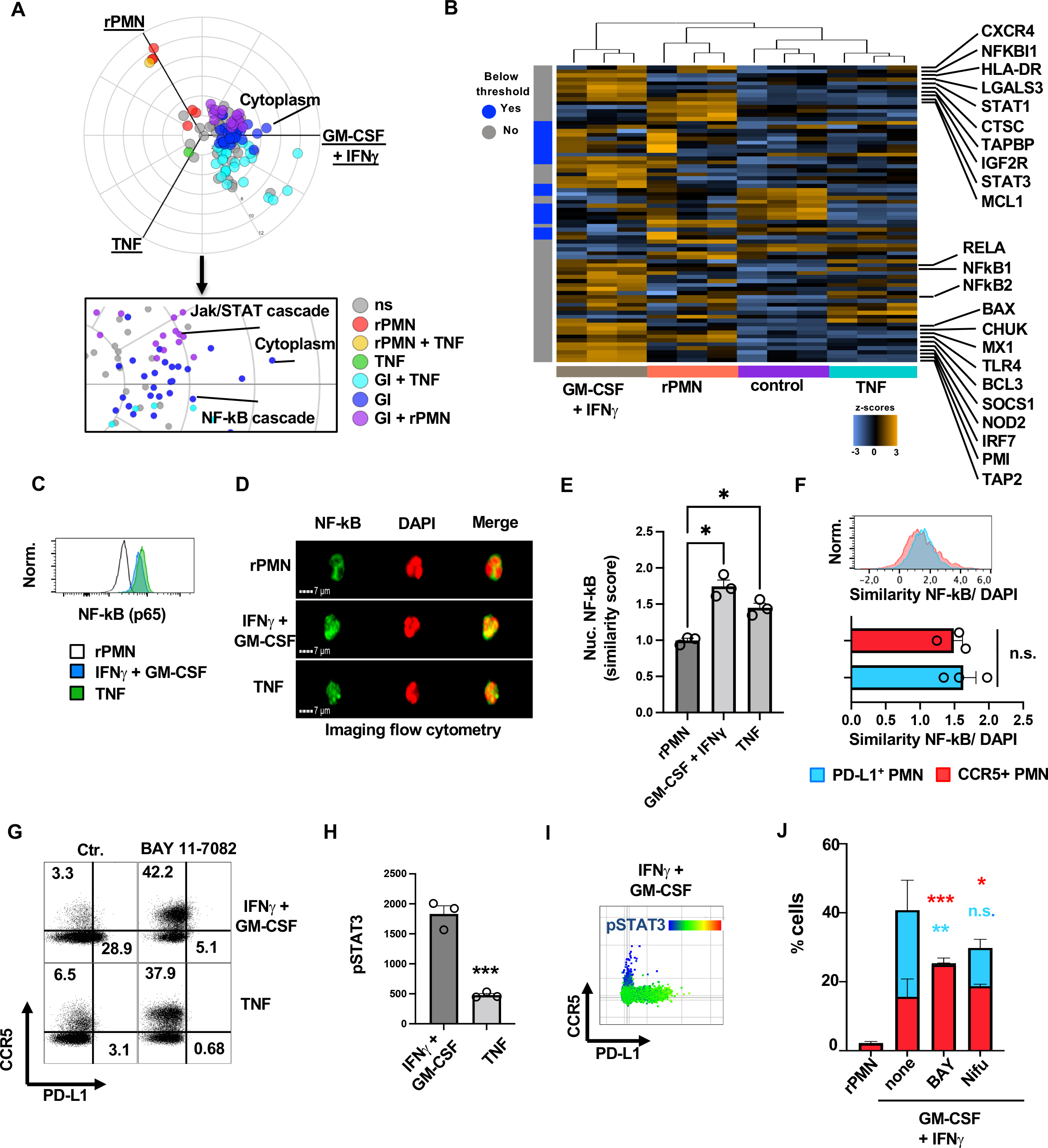
NF-kB and STAT3 promote PMN diversification towards PD-L1^+^ PMNs. **A**) A three-axis polar plot of gene signatures (GSs) derived from resting PMNs or primed PMNs (GM-CSF + IFNγ or TNF). Modules are color-coded based on statistical significance (false discovery rate < 0.05) for GS usage in different conditions. The box represents a zoomed-in view of the three-axis polar plot. The interactive version of this plot is available in the Supplementary material. **B**) Heatmap displaying normalized data from the GS “Cytoplasm,” with gene expression levels scaled to give all genes the same variance. Unsupervised clustering was used to generate the heatmap. Orange color represents high expression, while blue color represents low expression. This heatmap provides an overview of the data. **C**) Flow cytometric analysis of NF-kB expression. **D-E**) Imaging flow cytometric analysis of nuclear translocation of NF-kB in resting PMNs and after priming, as indicated. Representative images from three separate experiments (D) were included in a statistical analysis to calculate nuclear translocation using the similarity score (E, n=3, SEM, ANOVA, *p<0.05). **F**) Similarity score calculated using the rescaled Pearson correlation coefficient for CCR5^+^ and PD-L1^+^ PMNs after GM-CSF + IFNγ priming. The lower part of the panel shows quantification (n=3, SEM, t-test, n.s. = not significant). **G**) Flow cytometric analysis of CCR5 and PD-L1 expression on PMNs primed with either GM-CSF + IFNγ or TNF in the presence and absence of BAY 11-7082. **H**) Quantification of STAT3 phosphorylation after TNF or GM-CSF + IFNγ priming, presented as mean fluorescence intensity (MFI) obtained during flow cytometry (n=3, SEM, Mann Whitney U test, ***p<0.001). **I**) Distribution of pSTAT3 in CCR5^+^ or PD-L1^+^ PMNs after GM-CSF + IFNγ priming. **J**) Quantification of the percentage distribution of CCR5+ (blue) and PD-L1^+^ PMNs (red) after priming with GM-CSF + IFNγ in the presence and absence of BAY and Nifu (n=4, SEM, ANOVA, n.s. = not significant, *p<0.05, **p<0.01, ***p<0.001).

Flow cytometry confirmed an increased protein expression of NF-kB in GM-CSF plus IFNγ and TNF-primed PMNs (**Fig. 4C**). We then assessed nuclear translocation of NF-kB by imaging flow cytometry and correlated NF-kB and DAPI signal intensities. This analysis showed that NF-kB was active in both priming conditions (**Fig. 4D and 4E**). Similarly, a comparable nuclear translocation of NF-kB was found in CCR5^+^ and PD-L1^+^ PMNs, suggesting the relevance of NF-kB for both PMNs subpopulations (**Fig. 4F**). To experimentally elicit the relevance of NF-kB for PMN diversification, we inhibited NF-kB pharmacologically using the IKK inhibitor BAY 11-7082 (2.5µM) (**Fig. 4G**). Unexpectedly, BAY 11-7082 inhibited PD-L1 expression but induced upregulation of CCR5.

The findings that nuclear translocation of NF-kB occurred in both CCR5^+^ PMNs and PD-L1^+^ PMNs but inhibition of NF-kB dampened PD-L1 expression, suggested that NF-kB was necessary but not sufficient to induce PD-L1 expression. It is known from other systems that STAT3 signaling downstream of the IFNG receptor is essential for robust PD-L1 expression (Antonangeli et al., 2020; Cheng et al., 2018; Zerdes et al., 2018). Indeed, STAT3 was upregulated in GM-CSF plus IFNγ-primed PMNs (**Fig. 4B**) and was more strongly phosphorylated compared to TNF-primed PMNs (**Fig. 4H**). Moreover, STAT3 was phosphorylated in PD-L1^+^ PMNs but not in CCR5^+^ PMNs (**Fig. 4I**). To further test the influence of NF-kB and STAT3 on PMN diversification, PMNs were primed with GM-CSF plus IFNγ and simultaneously exposed to either BAY 11-7082 or the STAT inhibitor Nifuroxazide (Nifu). Both inhibitors significantly reduced in the proportion of PD-L1^+^ PMNs after GM-CSF plus IFNγ priming, while a significant CCR5 upregulation was only observed after BAY 11-7082 treatment (**Fig. 4J**). Based on these data, we propose that NF-kB inhibited the development of CCR5^+^ PMNs and - together with STAT3 - promoted the development of PD-L1^+^ PMNs.

### NF-kB halts actin disassembly-dependent upregulation of CCR5

Surface expression of receptors requires protein biosynthesis and transport to the cell surface. Many neutrophil proteins, such as CCR5, are stored in granules of PMN and can be presented at the cell surface upon priming or stimulation (Neuenfeldt et al., 2022b). This requires dynamization of the actin cytoskeleton and the degradation of the cortical F-actin barrier (Jog et al., 2007). It was recently shown that the *LPL* gene, encoding an actin-reorganizing protein, is upregulated in myeloid cells of the synovia of arthritis patients (Lewis et al., 2019). We found that priming induced higher LPL protein expression in PMNs (**Fig. 5A**). To also gain insight into the function of LPL, we quantified phosphorylation at Ser5 (**Fig. 5B**) and detected more robust phosphorylation of LPL 2h after GM-CSF plus IFNγ priming. This remained increased after 24h and was sensitive to the NF-kB inhibitor BAY 11-7082. Even though a trend to similar kinetics was observed with TNF, LPL phosphorylation was not significantly increased after 24 h. A direct comparison of the phosphorylation index between CCR5^+^ and PD-L1^+^ PMN showed stronger LPL phosphorylation in the latter (**Fig. 5C**).

**Figure 5.**
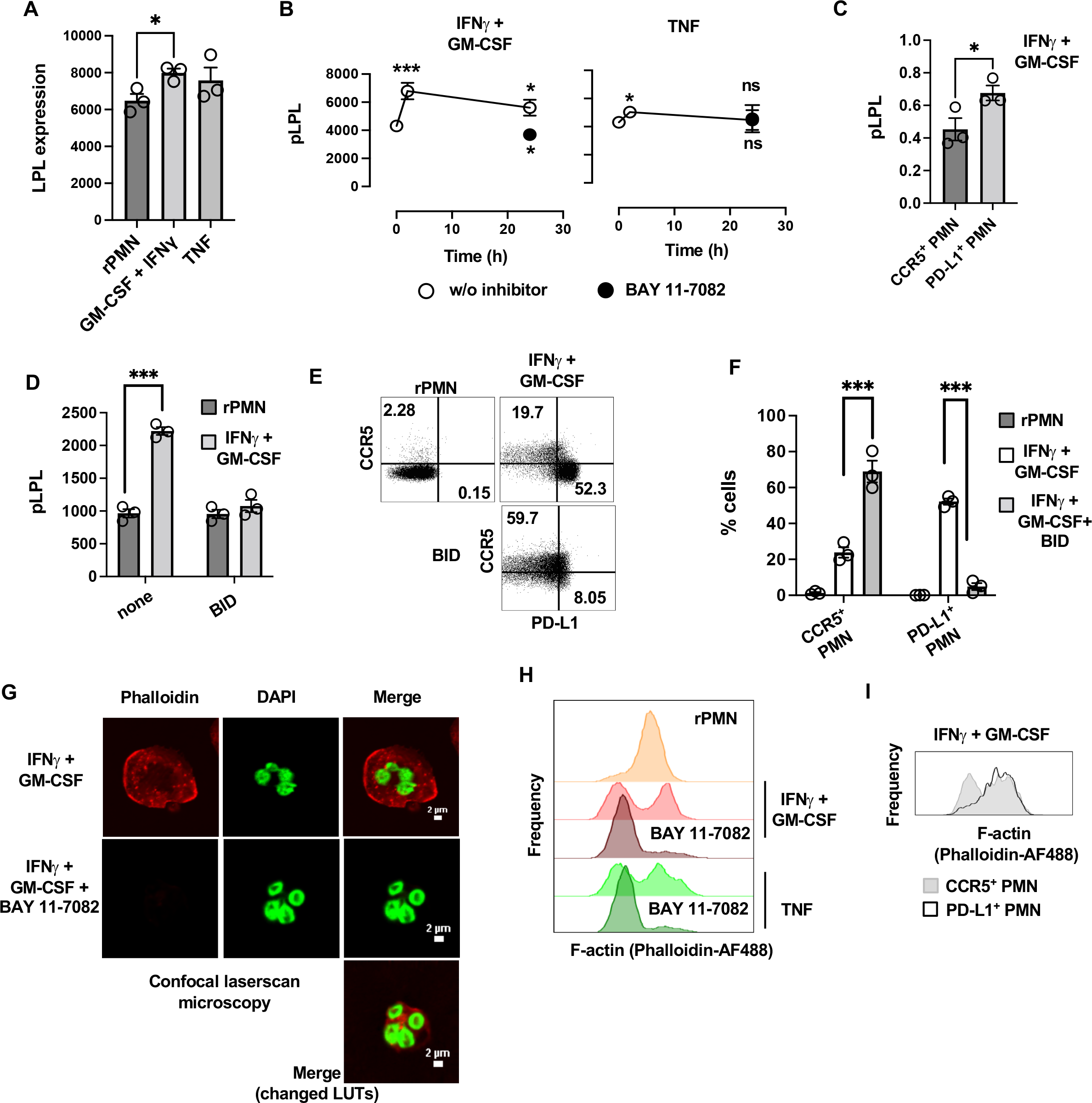
Actin dynamics modulate PMN diversification. **A**) Flow cytometric quantification of LPL expression (SEM, n=3, * p<0.05, ANOVA). **B**) Kinetics of LPL phosphorylation at Ser5. Phosphorylation was assessed using a phosphor-specific antibody and flow cytometry (SEM, n=3, *p<0.05, ***p<0.001, ANOVA, white circles = statistical comparison to rPMNs, black circles = statistical comparison to t=24 h in the absence of BAY 11-7082). **C**) Flow cytometric quantification of LPL phosphorylation index in CCR5^+^ or PD-L1^+^ PMNs (GM-CSF + IFNγ priming) (SEM, n=3, * p<0.05, t-test). **D**) Quantification of LPL phosphorylation in the presence or absence of BI-D1879 (BID) (SEM, n=3, ***p<0.001, ANOVA). **E-F**) Flow cytometric analysis of cytokine-induced PMN diversification towards CCR5^+^ or PD-L1^+^ PMNs in the presence or absence of BID. The numbers in the dot plots represent the respective percentage of cells (F, n=3, SEM, ***p<0.001, ANOVA). **G**) Confocal laser-scanning microscopic images of primed PMNs in the presence and absence of BAY 11-7082. **H**) Staining of F-actin (using phalloidin-AF488) in resting PMNs immediately after purification or after incubation with cytokines in the presence or absence of BAY 11-7082 for 24 h. **I**) Staining of F-actin (Phalloidin-AF488) in CCR5^+^ PMNs and PD-L1^+^ PMNs after GM-CSF plus IFNγ priming for 24 h.

To investigate the function of LPL phosphorylation for PMN diversification, we interfered with the corresponding signaling pathway using the LPL kinase inhibitor BI-D1879 (BID) (Wabnitz et al., 2021). This completely blocked LPL phosphorylation in GM-CSF plus IFNγ-primed cells (**Fig. 5D**). At the same time, the formation of PD-L1^+^ PMNs was inhibited while the number of CCR5^+^ PMNs increased significantly (**Fig. 5E and F**).

We have previously shown that inhibition of LPL phosphorylation leads to changes in the activation-induced actin reorganization and an overall decrease in F-actin content in T cells (Wabnitz et al., 2021). Actin reorganization can be determined by measuring F-actin using fluorescently labeled phalloidin. Confocal laser-scanning microscopy showed that the F-actin content of GM-CSF plus IFNγ-primed PMNs was lowered by BAY 11-7082 treatment (**Fig. 5G**). This finding was confirmed by flow cytometry for GM-CSF plus IFNγ and TNF priming where two BAY 11-7082-sensitive F-actin peaks were seen (**Fig. 5H**). These distinct F-actin^high^ and F-actin^low^ populations were typical for CCR5^+^ PMNs while PD-L1^+^ PMNs were homogenously F-actin^high^ (**Fig. 5I**). Note that physical stress induced actin-disassembly and an increase of CCR5 expression (**Fig. S7A and B**). Based on these data, we propose that actin-disassembly induced by stress or priming favors surface transport of granule-stored CCR5. NF-kB regulates LPL expression. LPL activation leads to a stabilization of the actin cytoskeleton ultimately leading to inhibition of CCR5 surface expression. This assumption is supported by Western blots showing a constitutive expression of CCR5 in whole cell lysates (**Fig. S7C**). Additionally, significant amounts of PD-L1 were detected exclusively in whole cell lysates obtained from GM-CSF plus IFNγ-primed PMN.

### Differential priming determines the functional outcome of PMNs

Next, the functional properties of PMNs subpopulations were examined. Since sorting of primed PMNs-induced cell death, we analyzed functional features in bulk primed PMNs combined with flow or imaging flow cytometry gating. First, primed PMNs were stimulated with PMA, ionomycin, or fMLP, and the degranulation capacity of specific and gelatinase granules was measured by increased expression of CD66b at the cell surface (**Fig. 6A**) (McLeish et al., 2013). For all three types of stimulation, significantly weaker exocytosis was observed in TNF-primed PMNs compared with GM-CSF plus IFNγ-primed cells. Residual exocytosis activity could be detected for TNF-primed PMNs after ionomycin or fMLP stimulation. In contrast, exocytosis of GM-CSF plus IFNγ-primed PMNs was stronger after ionomycin, and equally strong after fMLP stimulation compared with rPMNs. A weaker exocytosis activity was observed after PMA stimulation, which could be explained by NETosis induction.

**Figure 6.**
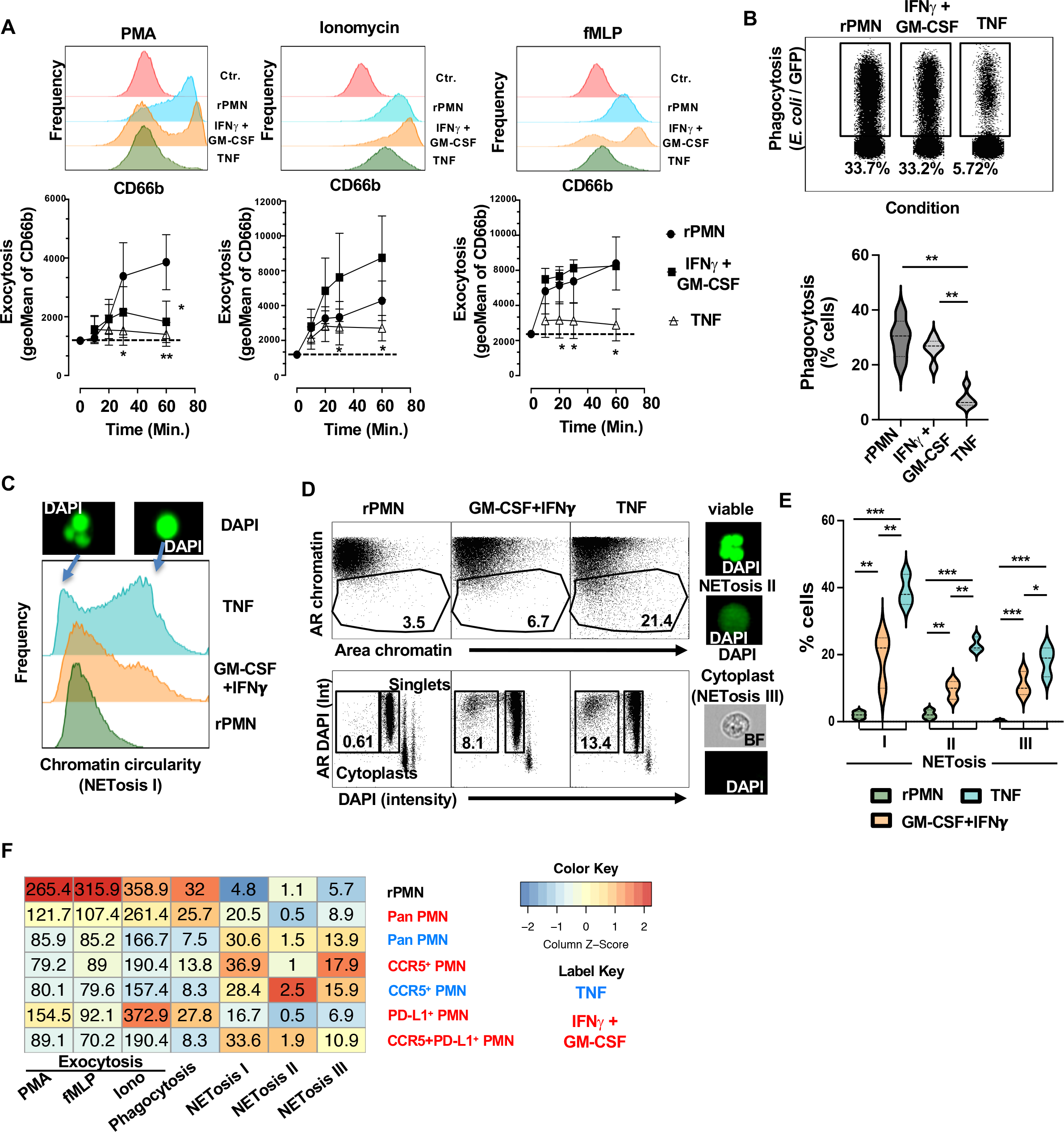
The dichotomous function of diversified PMNs. **A**) PMNs were stimulated as indicated for 60 minutes (upper part). A time course from 5 to 60 minutes is shown in the lower part. Exocytosis of granules was determined by measuring the mean fluorescence intensity (MFI) of CD66b surface expression using flow cytometry (SEM, n=3, *p<0.05, **p<0.01). **B**) Merged flow cytometry plot and percentages of *E. coli*-positive (GFP^+^) PMNs. The conditions are indicated above, and the percentages refer to the percentage within each condition. The figure is representative of 5 independent experiments, and a corresponding quantification is provided (SEM, n=5, **p<0.01). **C**) Nuclear morphology was calculated using the circularity feature (Circularity_Morphology (M07, Ch07)). Histograms show the nuclear morphology along the x-axis, and representative images of DAPI-stained cells are displayed. The figure is representative of 5 independent experiments. Quantification is shown in E. **D**) PMNs undergoing NETosis were determined based on nuclear morphology aspect ratio (y-axis; AR chromatin) and chromatin area (x-axis, area chromatin) using DAPI staining. Example images from the gates are shown on the right. In the lower part of the figure, quantification of one NETosis end-stage, namely the appearance of nucleus-less cytoplasts, is presented. The percentages refer to singlets, while doublets and triplets are shown for completeness. Quantification is shown in E. **E**) Violin plot displaying quantification and statistical analysis of the data presented in C and D. The plot includes data from 5 independent experiments and was statistically analyzed using ANOVA (n=5, *p<0.05, **p<0.01, ***p<0.001). **F**) Heatmap showing the assessment of the indicated PMN functions according to stimulation and PMN subpopulations.

Phagocytic ability of the cells was tested by incubation with GFP-labeled *E. coli* for 45 min. Ingested bacteria were subsequently determined by flow cytometry based on the green fluorescence signal present in the PMNs. Phagocytosis was significantly lower in TNF-primed PMNs than in PMNs primed with GM-CSF plus IFNγ or in rPMNs (**Fig. 6B**). These results are in line with the finding that TNF-primed PMNs contained lower TLR4 mRNA levels (**compare Fig. 4B**).

We have recently developed a method to quantify NETosis in suspension cells by quantifying the chromatin morphology using imaging flow cytometry (Neuenfeldt et al., 2022b). At a very early stage of NETosis, the nuclear morphology changes from lobed shape to round shape. Calculating the circularity of the nucleus showed hardly any round nuclei in rPMNs. Priming involving IFNγ resulted in a higher circularity score and a subpopulation with round nuclei. TNF-dependent priming resulted in even more cells with a high circularity score, i.e. round nuclei (**Fig. 6C and E**). In the progression of NETosis, nuclear dissolution and chromatin expansion occur, which can be quantified by increased area of chromatin (**Fig. 6D and E**). Again, more TNF-primed cells showed this NETosis phenotype compared to rPMNs or IFNγ plus GM-CSF-primed cells. A similar situation was seen in the appearance of cytoplasts detectable as chromatin-free “ghost” cells, which were more frequent in TNF-primed PMNs.

The difference in differential antibacterial capabilities of the cells may have resulted from diversification. Indeed, CCR5^+^ and CCR5^+^PD-L1^+^ double positive PMNs showed weaker exocytosis and phagocytosis with concomitant increased NETosis compared with rPMNs (**Fig. 6F**). In contrast, PD-L1^+^ PMNs showed a phenotype similar to rPMNs in exocytosis, phagocytosis, and NETosis.

### The type of PMN priming has differential effects on T cell immunity

To scrutinize the relevance of PMN priming for T cells, PMNs were loaded with the superantigen SEB, which bound mainly on HLA-DR positive PMNs (**Fig. 7A, upper part**). Autologous T cells formed few conjugates with these cells (**Fig. 7A, lower part**). CD69 was upregulated on T cells co-cultured with TNF-primed PMNs, but not with GM-CSF + IFNγ primed PMNs (**Fig. 7A, lower part**).

**Figure 7.**
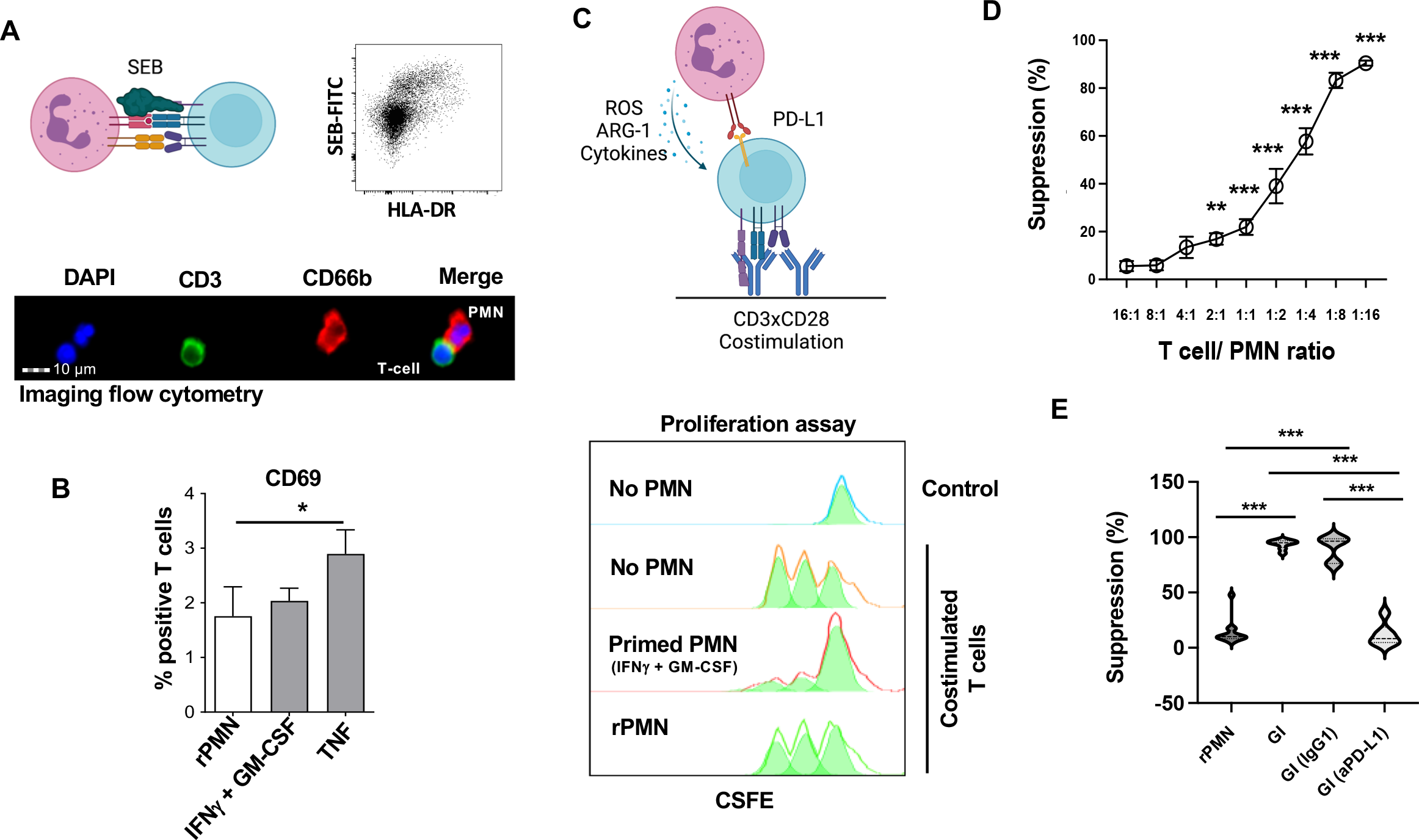
PD-L1-mediated inhibition of T cell proliferation. **A**) The upper part shows a schematic of T cell/PMNs interaction and stimulation of T cells via SEB (left). The binding of FITC-labeled SEB on HLA-DR^+^ cells is shown on the right. The lower part depicts imaging flow cytometry data of a T cell/PMN couple. **B**) The bar graph displays quantification of the activation marker CD69 expressed on T cells. The geometric mean of 5 independent experiments is shown (SEM, n=5, *p<0.05). **C**) Diagram illustrating the potential modulation of T cell response using PMNs as bystander cells. T cells (blue) are stimulated via cross-linked antibodies, while PMNs (purple) modulate T cell activation either indirectly via secreted substances or directly through receptor/ligand interactions. Cell proliferation of the indicated conditions is depicted as a CSFE dilution assay. **D**) T cell proliferation suppression assay showing the titration of the T cell/PMN ratio. PMNs were primed with GM-CSF + IFNγ. **E**) T cell proliferation suppression assay showing the influence of differently primed PMNs. PMNs (IFNγ + GM-CSF primed) were incubated with PD-L1 blocking antibody or an isotype control, followed by co-incubation with T cells as described in B (GI: GM-CSF + IFNγ-primed PMNs, GI (IgG): GM-CSF + IFNγ-primed PMNs in the presence of isotype control antibodies, GI (aPD-L1): GM-CSF + IFNγ-primed PMNs in the presence of PD-L1-blocking antibodies; n=3, t-test, ***p<0.001).

To determine whether GM-CSF + IFNγ - primed PMNs can attenuate T cell immune responses via PD-L1 expression, T cells were co-stimulated via plate-bound antibodies against CD3 and CD28 (polyclonal activation) and co-incubated with primed PMNs (**Fig. 7B**). T cell activation was measured by a 72h CSFE-proliferation assay to allow upregulation of PD1 on T cells. Costimulated T cells divided on average three times during the 72h period. T cell division was significantly reduced in the presence of GM-CSF + IFNγ primed PMNs starting at a T cell to PMN ratio of 2:1 (**Fig. 7C**). Incubation with rPMNs did not interfere with T cell proliferation. In addition, GM-CSF + IFNγ - primed PMNs inhibited co-stimulation-dependent T cell proliferation significantly stronger than TNF primed cells (**Figs. 7C and S8A**). The T cell suppressive effect of GM-CSF + IFNγ-primed PMNs could be abolished by PD-L1-blocking antibodies but not by an isotype control (**Fig. 7D**). Supernatants of rPMNs, primed PMNs (GM-CSF + IFNγ) or T cell / primed PMN (GM-CSF + IFNγ) co-cultures did not induce significant suppression of T cell proliferation (**Fig. S8B**). Therefore, GM-CSF + IFNγ-primed PMNs suppress T cells via PD1/PD-L1 interaction.

## Discussion

The notion that neutrophilic granulocytes (PMNs) are a homogeneous cell population has been challenged by recent studies, revealing significant diversification and plasticity among PMNs in different tissues and conditions in both mice and humans (Grieshaber-Bouyer et al., 2021; Ng et al., 2019; Silvestre-Roig et al., 2019; Xie et al., 2020). However, the molecular mechanisms underlying this diversification remained poorly understood. Furthermore, there is ongoing debate regarding whether mature PMNs can be reprogrammed by the local microenvironment (Ballesteros et al., 2020; Grieshaber-Bouyer et al., 2021; Rosales, 2018; Xie et al., 2020).

In this study, we uncovered the remodeling of the actin cytoskeleton and the combined activity of NF-kB and STAT3 as triggers for diversification of mature PMNs in inflammatory settings. We primed PMNs *in vitro* with GM-CSF and IFNγ, which are known to prolong PMNs’ survival and induce surface transport of granule-stored CD66b (Kamp et al., 2013; Neuenfeldt et al., 2022a). We observed that these cytokines prevented PMN apoptosis, as indicated by the low number of Annexin-V/7-AAD positive cells (Balta et al., 2017). It is worth noting that viable cells can also expose phosphatidylserine (PS) and become Annexin-V positive during activation, without undergoing DNA fragmentation or displaying other apoptotic signs (Dias-Baruffi et al., 2003; Fischer et al., 2006; Frasch et al., 2004; Segawa et al., 2011; Stowell et al., 2009). Consequently, we employed various approaches to assess apoptosis in our current *in vitro* study. We did not observe extended DNA fragmentation in cytokine-primed cells analyzed by imaging flow cytometry. Similarly, utilization of the live/dead stain amine-binding dye Zombie and Apotracker (which binds to PS independent of calcium avoiding staining-related artifacts on PMN function) did not show increased apoptosis after 24 hours. However, it increased markedly after three days of incubation. Furthermore, in our preliminary experiments, we observed no discernible rise in apoptosis among the PMNs *ex vivo* from samples obtained from patients during the initial 5-hour period. However, after the 4-hour mark, we noticed changes in the phenotype of the neutrophils, as e.g. upregulation of CCR5. To ensure reliable results, we implemented a protocol where patient samples would only be utilized if they reached the laboratory within two hours. Overall, our control experiments confirmed that no significant alterations in the PMN phenotype occurred under these pre-analytical conditions and our *in vitro* priming protocols allowed us to analyze PMN diversification paths while minimizing apoptosis.

Our diversification model provides mechanistic insights into how PMNs integrate their immediate environment and translate it into effector functions. The emergence of CCR5^+^ PMNs, known to contribute to accelerated NETosis, was driven by actin-remodeling-induced upregulation of CCR5 and TNFR2 (Neuenfeldt et al., 2022a). Our observations were supported by a recent study showing TNFR2-mediated NETosis in neutrophils of mice infected with *S. aureus* (Youn et al., 2023). Factors such as aging and shear stress can modulate the actin cytoskeleton, and PMNs have the ability to mechanotransduce shear forces into functional adaptations and alter their surfaceome (Fine et al., 2016; Hilmo and Howard, 1987; Sheikh et al., 2003; Shive et al., 2000; Sun et al., 2021). CD62L, a receptor downregulated in response to shear forces, was also weakly expressed in CCR5^+^ PMNs (this work and (Neuenfeldt et al., 2022a)). The modulation of the actin cytoskeleton and exocytosis of receptors offers energetically efficient mechanisms for PMNs to quickly respond to environmental changes on-demand, without relying on gene regulation processes including transcription, translation, and surface transport. The development of CCR5^+^ PMNs can be considered a canonical pathway triggered by the disassembly of the actin cytoskeleton. Accordingly, these cells occurred under various pro- and anti-inflammatory conditions, as e.g. IL-10 or TNF. The low actin dynamics may also explain the functional properties of CCR5^+^ PMNs, such as reduced phagocytosis and enhanced NETosis, as phagocytosis requires actin cytoskeleton remodeling to form the phagocytic cup, and actin depolymerization is an early event during (and promoting) NETosis (May and Machesky, 2001; Thiam et al., 2020).

Activation of NF-kB by factors present in the microenvironment prevents apoptosis in PMNs, stimulates their effector responses, and influences actin dynamics (Khoyratty et al., 2021; Marion et al., 2012; Pillar et al., 2019; Stanisavljevic et al., 2011). In our research, we have discovered that the actin-bundling protein LPL plays a functional role in connecting NF-kB to the actin cytoskeleton, resulting in the stabilization of F-actin. This NF-kB-LPL pathway hinders the typical development of CCR5^+^ PMNs. Through a combination of stochastic mathematical modeling and experiments, Czerkies et al. demonstrated marked differences in the kinetics of NF-kB activation between Poly(I:C) and LPS stimuli (Czerkies et al., 2018). Consistent with these findings, our results show that NF-kB activation opposes CCR5 expression, and Poly(I:C) challenge leads to development of PD-L1^+^ PMNs at the expense of CCR5^+^ PMNs in mice. The upregulation of the immune checkpoint receptor PD-L1 is dependent on the activation of STAT3 in conjunction with NF-kB. This activation can occur through various factors, such as IFNγ or LPS. The concurrent activation of STAT3 and NF-kB is adequate for the generation of PD-L1^+^ PMNs. Consequently, in a cytokine mixture, the presence of a single cytokine that induces significant STAT3 activation may be sufficient for PD-L1 expression. Hence, PD-L1^+^ PMNs are observed in the combination of GM-CSF, IFNγ, and TNF, although IFNγ alone is capable of inducing PD-L1 expression. Notably, activation of NF-kB by the other cytokines tends to exhibit a synergistic effect and the anti-inflammatory *in vitro* regime using IL-10 did not induce PD-L1 expression on PMNs. Functionally, PD-L1^+^ PMNs shift towards an immunosuppressive phenotype, as evidenced in tumor settings, experimental bacterial sepsis, and thermal injury scenarios (de Kleijn et al., 2013; Qi et al., 2021; Thanabalasuriar et al., 2021; Wang et al., 2017; Wang et al., 2021).

Although PMNs are primarily associated with bacterial defense, they also play crucial roles during viral infections (Bowers et al., 2014; Tang et al., 2019). In our investigation of peripheral blood from severe COVID-19 patients, we observed PMN diversification with increased expression of both PD-L1^+^ and double positive (CCR5^+^/PD-L1^+^, DP) PMNs, which might be the consequence of a cytokine storm, typical for severe COVID-19 patients. Consequently, occurrence of PD-L1^+^ PMNs is rather associated with disease severity (MacDonald et al., 2021; Rice et al., 2023). This could be due to their immunosuppressive activity or the finding that DP PMN showed higher NETosis activity. The latter is associated with immunthrombosis (Cesta et al., 2023; Skendros et al., 2020; Zuo et al., 2020). Notably, CD66b upregulation and the occurrence of DP PMN was even maintained in convalescent patients, 3 months after recovery. This result represents one of the first descriptions of a “immunological scar” manifesting in PMNs and such a long-lasting PMNs phenotype alteration might have pathological relevance, since DP PMNs showed higher NETosis activity which could contribute to immune thrombotic dysregulation (Nicolai et al., 2020; Patell et al., 2020; Zuo et al., 2020).

To gain insights into PMNs in cases of low-grade inflammation, we examined PMNs in patients with osteoarthritis. Osteoarthritis is a prevalent disease and a significant public health concern, particularly in aging populations of developed countries. It diminishes the quality of life due to joint pain and functional limitations. Initially, osteoarthritis was considered a degenerative and non-inflammatory joint disorder. However, it has now been established that low-level inflammatory processes sustain the disease and contribute to its progression (Kapoor et al., 2011; Scanzello, 2017). In our study, we analyzed samples from patients undergoing planned joint-replacement surgery. Hence, this cohort likely was characterized by chronic low-grade inflammation and a long prior disease course. Even in the absence of acute inflammation, we observed the presence of both PD-L1 and CCR5 expression on PMNs in the synovial fluid. CCR5^+^ and DP PMNs are prone to undergo NETosis, a process that releases nuclear components, autoantigens, and tissue-degrading enzymes such as proteases into the extracellular space (Gupta and Kaplan, 2016; Jorch and Kubes, 2017; Kessenbrock et al., 2009; Villanueva et al., 2011). Therefore, the existence of these cells in the synovial fluid of osteoarthritis patients could contribute to the persistent inflammation observed. Further studies correlating PMN diversification and subpopulation composition with disease progression in a larger cohort of osteoarthritis patients are necessary to validate this hypothesis.

Although many differences in gene expression, morphology and function exist between murine and human neutrophils, activation can induce a conserved core inflammation program (Hackert, 2022). In line with this, our present work further identifies conserved diversification paths leading to CCR5^+^ and PD-L1^+^ cell states. This leads to the appearance of similar subgroups during inflammations or infections. Since the number of PMN subgroups depends on the micro-milieu, PMN diversification paths may also contribute to the course of a disease. Furthermore, these findings have implications for patient treatment. In this regard, we were able to demonstrate that the presence of CCR5^+^ PMNs is associated with resistance to anti-TNF treatment in patients with colitis (Neuenfeldt et al., 2022a). It should be noted, that our approaches were focused on PD-L1 and CCR5 expression and may therefore have led to an underestimation of the diversification potential of PMNs. However, this underestimation can also be seen as a strength of the study, because it was possible to identify dichotomous PMNs development using small flow cytometry-based panels and simple gating strategies, paving a simplified way for future diagnostic procedures.

## Author contribution

N.H., J.C.S and G.W. conceived the idea for the manuscript and drafted the initial manuscript, N.H., J.C.S, T.E. and G.W. implemented the R code and supported algorithms, G.W. developed overall research goals, S.A. designed and supervised mice experiments, N.H., J.C.S, R.G.B., S.A. and T.E. data curation, guided data analysis, interpretation and validation, H.P., U.M., U.D., and T.R. managed and interpreted patient samples, Y.S. provided resources, N.H., J.C.S, F.S.N., F.K. and J.H. performed and analyzed experiments, G.W. conceptualized and designed the study, supervised the entire project and wrote the manuscript.

## Acknowledgements

The authors thank Thomas Giese for help with COVID-19 patients’ blood, Ulrike Dapunt for help with osteoarthritis patient samples, Manina Günter for technical assistance and Inaam Nakchbandi for critical reading of the manuscript.

## Material and Methods

### Reagents

The following reagents were used in the study: IFNγ (10 ng/µl, R&D systems), G-CSF (5 ng/ml, R&D systems), GM-CSF (100 U/µl, R&D systems), BAY11-7082 (2.5 µM, Merck Sigma-Aldrich), LPS (100 µg/ml), IL-8 (10 ng/ml, R&D systems), IL-10 (200 U/ml, R&D systems), fMLP (100 µM, R&D systems), LPS (100 µg/ml, R&D systems), fMLP (100 nM), TNF (0.1 ng/µl, R&D systems), IL-10 (200 U/ml, R&D systems), GM-CSF (1000 U/µl, R&D Systems), *Staphylococcus*-Enterotoxin-B (SEB) (5 µg/ml, Merck Sigma-Aldrich), DAPI (Merck Sigma-Aldrich), BI-D1870 (5 µM, BML-EI457 Enzo), Nifuroxazide (25 µM, Merck Sigma-Aldrich). The antibodies for detection of LPL and pLPL were generated in the laboratory (Wabnitz et al., 2021).

### Blood donation, isolation of PMNs and T cells and PMN priming

Blood was voluntarily collected from healthy donors by peripheral venipuncture. Exclusion criteria for healthy donors were the presence of an acute infection or a recent surgical procedure. The study was approved by the ethics committee of the University of Heidelberg (IRB approved ethics vote S-285/2015) and was conducted following the principles of the Declaration of Helsinki. Written informed consent was obtained in advance.

PMNs were isolated from 30-90 ml of peripheral blood obtained from healthy donors. The isolation process involved density centrifugation. Specifically, 30 ml of whole blood was combined with 20 ml of PolymorphPrep™ and centrifuged at 535 g for 35 minutes. The layer containing PMNs was collected, washed with a 0.45% sodium chloride solution in a 1:1 ratio, and centrifuged at 443 g for 10 minutes. To eliminate erythrocyte contamination, the resulting pellet was mixed with 20 ml of a 0.2% sodium chloride solution. Hemolysis was then halted after 10 seconds by adding 20 ml of a 1.6% sodium chloride solution. After two additional washes, the pellet was resuspended in RPMI + 10% FCS. Pipetting was performed with wide lumen pipettes and reduced to a minimum to avoid shear stress.

Isolation of peripheral blood mononuclear cells (PBMCs) was performed via density centrifugation through a sucrose-epichlorohydrin copolymer (FicoLite). T cells were purified from PBMC by magnetic-bead-based negative T-cell isolation (Miltenyi Biotec, Bergisch Gladbach, Germany) as described before (Wabnitz et al., 2010b). PMNs were obtained from the precipitated cells by lysing the erythrocytes with ACK buffer. The purified PMNs were cultured at a density of 1 × 10^6^ cells/ml at 37 °C and 5% CO2 in 6-well plates (IL-8: 10 ng/ml, IL-10: 200 U/ml, TNF: 0.1 ng/ml, G-CSF: 5 ng/ml, GM-CSF: 100 U/ml, IFNγ: 10 ng/ml, LPS: 100 µg/ml, fMLP: 100 nM). In some experiments cells were incubated in the presence of BAY 11-7082 (Sigma, B5556).

### Patients’ samples

In pre-experiments, PMNs of blood samples were analyzed hourly for 5 h. PMNs started to change CCR5 expression after 4 hours. Patient samples were therefore only used if they arrived at the laboratory within 2 hours, which was possible with the short distances between facilities in Heidelberg. According to our control experiments, no significant PMN phenotype occurred under these pre-analytical conditions.

Patients with osteoarthritis were selected for whom surgical intervention for joint replacement was imminent at the “Center for Orthopaedics and Traumatology” at the Heidelberg UniversityHospital. After obtaining informed consent (IRB-approved ethics vote S-119/2017), both synovial fluid and peripheral blood were obtained as part of the scheduled procedure. The joint was punctured directly before incision. Blood sampling was also performed before the start of surgery under anaesthesia. The samples were shipped by courier to the laboratory for immediate processing. The synovial fluid was transferred to a heparinized 50 ml syringe and mixed with culture medium at a ratio of 1:3. The mixture was cleared using a 70 µm cell strainer, washed twice with culture medium, centrifuged at 300g for 10 minutes, and collected at a concentration of 10^6^ cells/ml in culture medium. Heparin anti-coagulated blood samples of severe COVID-19 were collected during their ICU stay (IRB approved ethics vote S-148/2020) and phenotyping was performed within 24h. For whole blood staining, 200 µl of blood was mixed with antibodies and incubated for 30 min at 4°C in the dark. Erythrocytes were lysed with BD FACS^TM^ Lysing Solution before measuring the cells with the flow cytometer.

### Flow cytometry

Flow cytometry was performed on a BD LSRII flow cytometer and by using the following antibodies: CD66b PerCP-Cy5.5, BioLegend 305108; CCR5 PE Miltenyi 130-106-223; PD-L1 APC, BD Biosciences 563741; PD-L2 PE-Vio770 Miltenyi 130-105-828; ICOSL BV421 Biosciences 564276; CD80 BV650 BioLegend 305227; CD15 BV605 Biosciences 562980; CD49d PE-Cy5 BD Biosciences 559880; CD58 PE-Cy5 BioLegend 330909; CD83 PE-Cy7 BD Biosciences 561132; CD86 PE-CF594 BD Biosciences 562390; CD209 AF647 BioLegend 330112; CXCR1 BioLegend 320612; HLA-DR AF700 BioLegend 307625. Where indicated T-cells and monocytes (CD3, CD14) cells were excluded. Data were exported from the FACSDiva software as FCS 3.0 files and gated on CD66b positive, zombie negative PMNs using FlowJo (v10.8.1). Where indicated, gated events and their median fluorescence intensity values were concatenated, exported and imported as CYT-object into CytoTree (v1.4.0) and R (v. 4.1.2). The dataset was log-transformed. FlowSOM clustering, dimensionality reduction and tree-shaped trajectories were calculated using the algorithms implemented in the CytoTree package and by using the packages flowCore (v2.6.0) and destiny (v3.8.1). For visualization ggplot2 (v3.3.6), viridis (v0.6.2) and RColorBrewer (v1.1-3) were applied.

### Imaging flow cytometry

PMNs were subjected to analysis by imaging flow cytometry (IFC) for their formation of neutrophil extracellular traps (NETs) or nuclear translocation of NF-kB as described before (Neuenfeldt et al., 2022b). Briefly, 2×10^6^ cells were pelleted at 300 g and fixed with 1.5% PFA solution. Fixed PMNs were incubated with 100 µl of antibody mix for 40 minutes in the dark for staining of the membrane-bound target proteins. In order to stain of the nucleus with DAPI a cell permeabilization step was performed using 0.1% saponin. After this staining procedure, the cell images were immediately acquired using the ImageStream^®^ IsX MkII IFC and analyzed in the IDEAS analysis software. To evaluate the formation of NETs, 3 different stages of cell death were recorded. The roundness of the nuclei was determined by calculating a circularity index, the size (Area feature) and sharpness of the nuclei (Gradient RMS) were used to detect cells with swollen nuclei, and the percentage of nucleus-less cytoplasts was determined as DAPI negative CD15 positive events. Nuclear fragmentation was determined by calculating the area of 40% thresholded image of DAPI-stain. The Amnis^®^ NF-kB Translocation Kit (Luminex, ACS100000) and an NF-kB (p50) antibody (Luminex 4700-1674) were used to measure the nuclear translocation of NF-kB according to the manufacturer’s instruction.

### Gene set and pathway analysis

RNA was extracted from both rPMN and primed PMN samples obtained from three healthy blood donors. The isolation of PMN involved gradient centrifugation followed by magnetic bead-based separation. Total RNA was then purified from 10,000 cells using the MagNA Pure system (Promega, Mannheim, Germany) at our in-house molecular immunodiagnostic facility, following the manufacturer’s instructions. The quantification of all RNA samples was performed using the Qubit RNA assay kit (Thermo Fisher Scientific, Waltham, Massachusetts), and RNA integrity was evaluated using the Agilent 2100 Bioanalyzer system. Gene expression analysis was conducted in collaboration with the NanoString Core facility in Heidelberg. Briefly, 25 ng of total RNA (5 µL/sample) was combined with the nCounter® reporter CodeSet (3 µL) and nCounter® capture ProbeSet (2 µL) along with hybridization buffer (5 µL) for an overnight hybridization reaction at 65 °C. The reaction was then cooled to 4 °C, and the samples were purified, immobilized on a cartridge, and the data were assessed using the nCounter SPRINT Profiler and data were uploaded to GEO database. For a more comprehensive description of the sample preparation process, we refer to the original publication pertaining to these specific samples. (Neuenfeldt et al., 2022a). Here, corresponding RCC files of the dataset GEO: GSE194366 were imported into nSolver Analysis software (v4.0) and analysis of read-count data including QC, differential expression, normalization and thresholding (p-value threshold: 0.05) was performed. GSA, PCA and KEGG analysis were achieved using the plugin nCounter Advanced Analysis (v2.0.115). GSAs were exported as csv-files to produce radar plots with volcano3D (v2.0.8), volcano3Ddata (v0.1.0), and R (v. 4.1.2) (Lewis et al., 2019).

### T cell assays

PMNs were pelleted by centrifugation (300g, 5 min) and suspended in 50 µl culture medium containing 1.25 µg SEB/10^6^ cells. After 15 minutes of incubation at room temperature, 5 ml culture medium was added, and cells were spun down by centrifugation. 1×10^6^ PMNs were mixed with 3×10^6^ T cells, centrifuged at 400 g for 5 minutes and resuspended in 200 µl PMNs culture medium. The mixtures were incubated at 37 °C and 7.5% CO2 for 24 h. CD69-expression was analyzed by flow cytometry.

The proliferation of PBT was induced via CD3 antibody (20 ng/ml, clone: OKT3) and CD28 antibody (5 µg/ml, BD Biosciences 555725) bound to the surface of a 96-well Nunc MaxiSorp^TM^ microtiter plate. 3×10^6^ T cells were taken up in 300 µl of a 1 µM CFDA-SE-PBS solution in a 1.5 ml tube, mixed well and incubated for 15 minutes at 37 °C and 5% CO2. Subsequently, the cells were washed twice with 1 ml PBS and then picked up in 1 ml PMNs culture medium. Different numbers of primed PMNs were added as indicated and each experiment was performed in triplicate. In addition, six positive and six negative controls were included. T cells in wells coated with CD3 and CD28 antibodies served as positive controls, whereas T cells in uncoated wells were used as negative controls. Blocking experiments were performed with 10 µg of a PD-L1 blocking antibody (clone: MIH1) (de Kleijn et al., 2013). A sample set was incubated in parallel with an IgG1 isotype control antibody at the same concentration for negative controls.

### Phenotyping of spleen cells of mice

The experiments were performed with 7-week-old female C57BL/6JRj mice (Janvier, France). Animal procedures were carried out according to the protocol approved by the Regierungspräsidium Karlsruhe (Permit Numbers: G198/21 & G61/21). Mice were treated intravenously with 200 µl PBS. 28 days later, the mice were challenged intraperitoneally with 200 µl PBS or LPS from *S. enterica* serovar *thyphimurium* (20µg/mouse; Sigma), Pam3CSK4 (100 µg/mouse; InvivoGen), or Poly(I:C) (100 µg/mouse; InvivoGen) in 200 µl PBS for 24 h. Preparation of single cell suspensions from spleen was performed as previously described (Autenrieth et al., 2012). Total cell numbers were determined by trypan blue exclusion. Antibodies used are listed in Table S1. All washing and incubation steps were performed with cell staining buffer (BioLegend). Zombie NIR (BioLegend) was used to exclude dead cells according to the Manufacturer’s instructions. Blocking of Fc receptors was done before staining by incubating cells at 4 °C for 15 minutes with anti-FcγRII / III mAb. Extracellular staining was performed for 1 h at 4 °C. After washing cells were fixed and permeabilized using the True-Nuclear™ Transcription Factor Buffer Set (BioLegend) followed by intracellular staining for 30 minutes at room temperature. Samples were acquired on a spectral flow cytometer AURORA (Cytek Biosciences).

### Statistics and data presentation

For comparisons means t-test (two groups) or ANOVA test (three or more groups) with Fisher’s exact test as included in the software GraphPad Prism (v9.3.1) was used. Asterisks represent the following *p*-value ranges: *p* > 0.05, n.s.; *p* < 0.05, *; *p* < 0.01, **; *p* < 0.001, ***. If sample images or figures are depicted, they are representative for at least three repetitions. Models were created with BioRender.

## Supplement material

**Figure S1.**
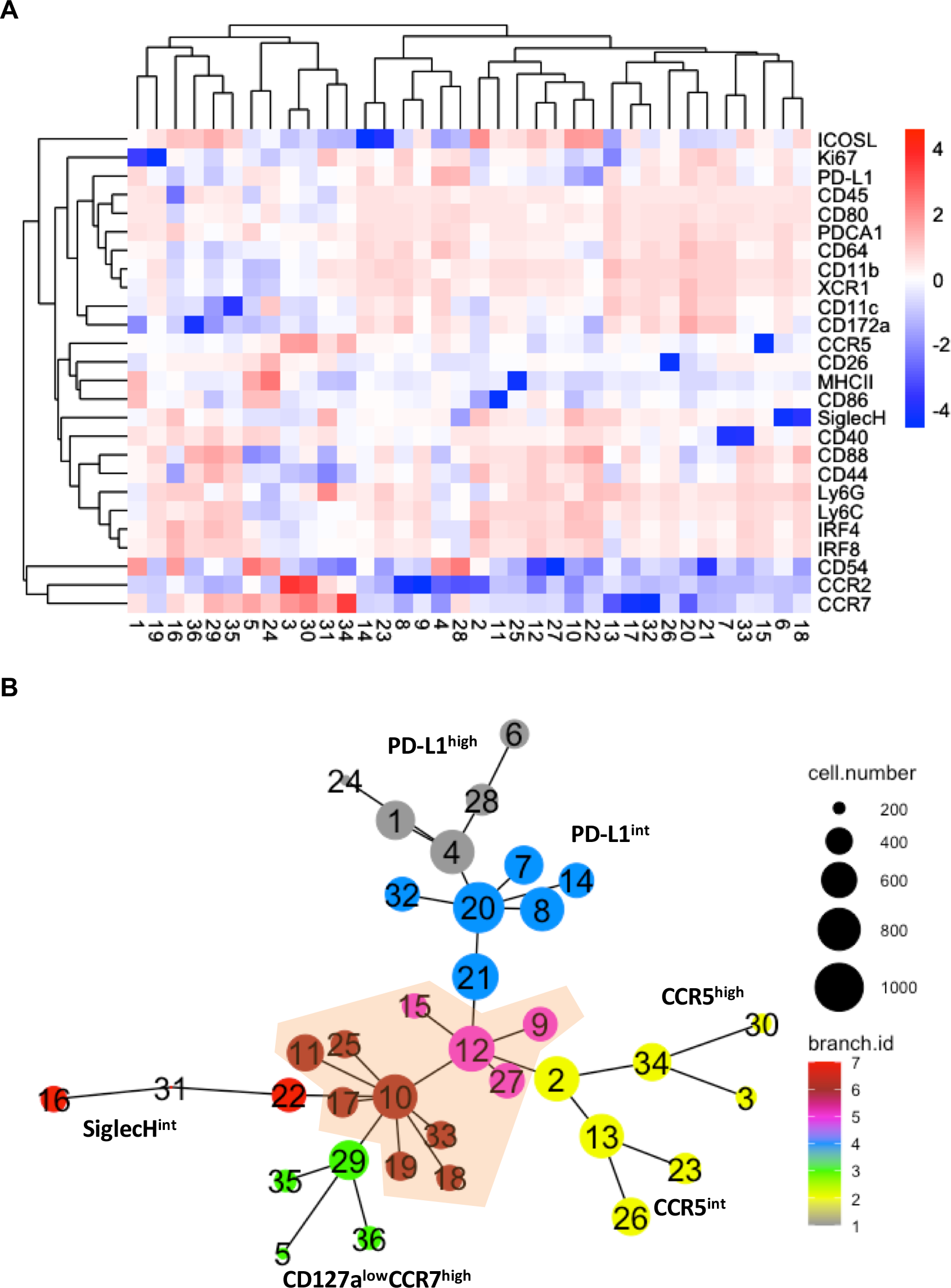
Expression analysis of PMNs from challenged mice. **A-B**) Self-Organizing Map (SOM) cluster heatmap (A) and tree representation (B) of spleen PMNs from challenged mice (compare Fig. 1). Lead expression marker annotation is included manually. **C**) Original flow cytometry dot plots showing CCR5 and PD-L1 expression on PMNs.

**Figure S2.**
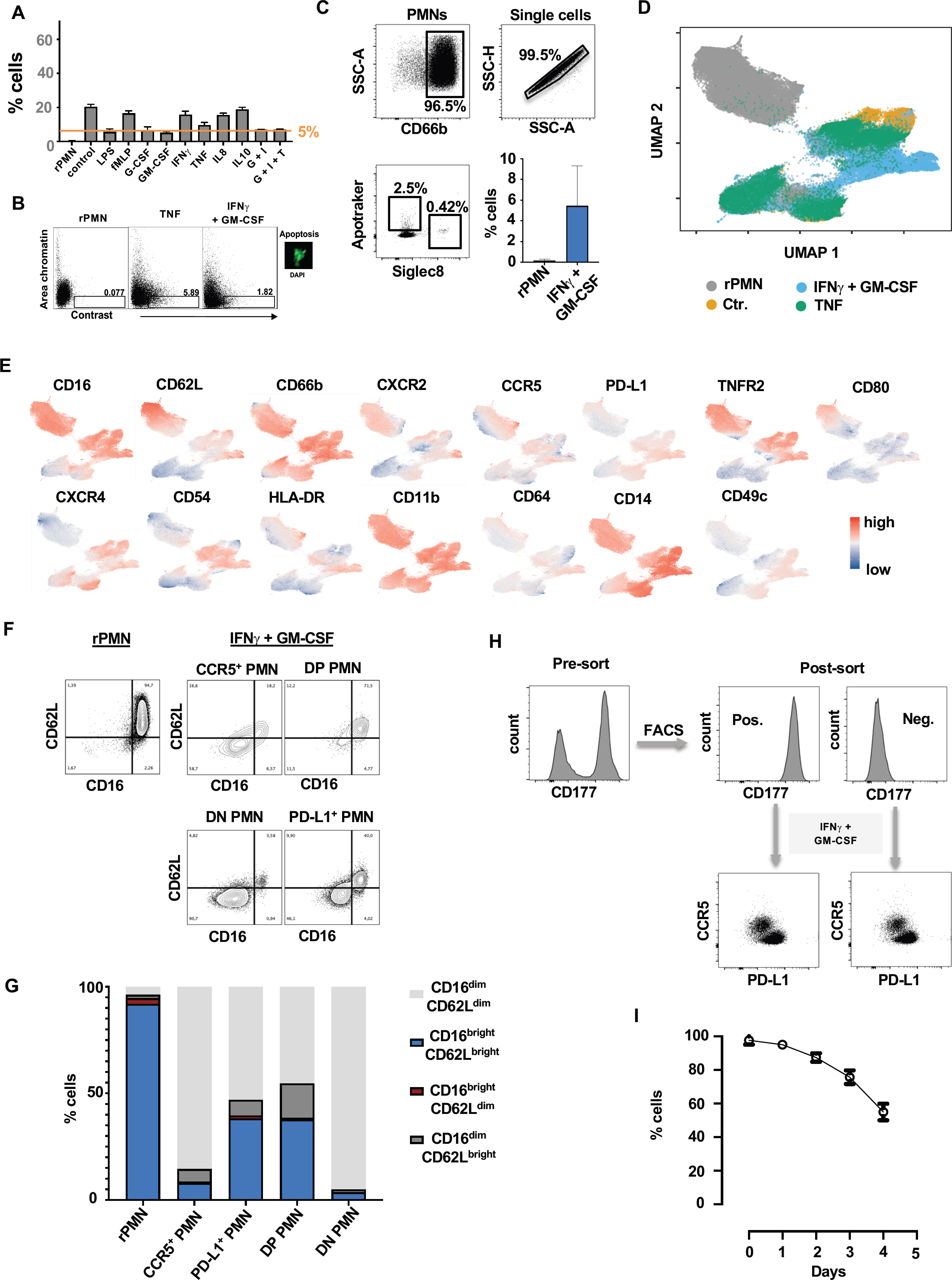
Survival and expression profile of *in vitro* primed human PMNs. **A**) Quantification of Zombie^+^ PMNs across conditions (n=3, SEM). **B**) Imaging flow cytometry analysis of apoptosis according to nuclear fragmentation. Low area of the chromatin staining combined with increased contrast is typical for apoptotic cells. The figure is representative of 5 independent experiments. **c**) Quantification of Apotracker^+^ resting PMNs and PMNs primed with GM-CSF + IFNγ for 24 h (n=3, SEM). **D-E**) UMAP projections of the indicated PMNs (C). The dataset is colored according to condition (C) or receptor expression (D). **F-G**) Flow cytometry dot plots showing the expression of CD16 and CD62L on resting PMNs or PMN subpopulations arising after priming with GM-CSF + IFNγ (E). The quantification represents the mean of 3 independent experiments. **H**) Resting PMNs were FACS-sorted based on CD177 expression. Both CD177^+^and CD177^+^ cells were primed with GM-CSF + IFNγ, and the expression of CCR5 and PD-L1 was assessed after 24 h. **I**) Measurement of PMN survival using Zombie staining upon priming with GM-CSF + IFNγ for 4 days (n=3, SEM).

**Figure S3.**
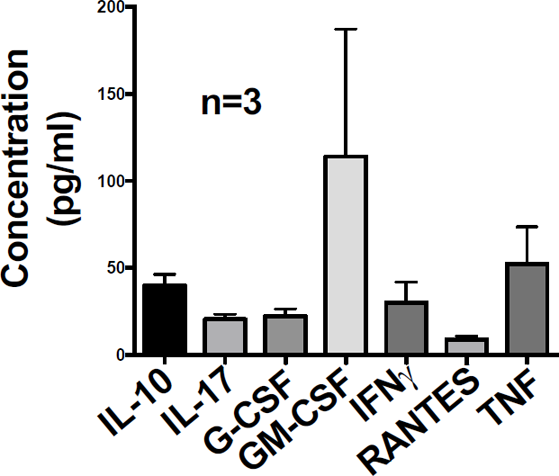
Cytokine profile in OA joints. Human Cytokine/Chemokine Magnetic Bead Multiplex Assay was used to measure cytokines in 3 different synovial fluid samples from patients with OA. The assay allowed the simultaneous analysis of multiple cytokine and chemokine biomarkers using bead-based and Luminex technology. The array was performed according to the manufacturer’s instructions (Merck KGaA, Darmstadt, Germany).

**Figure S4.**
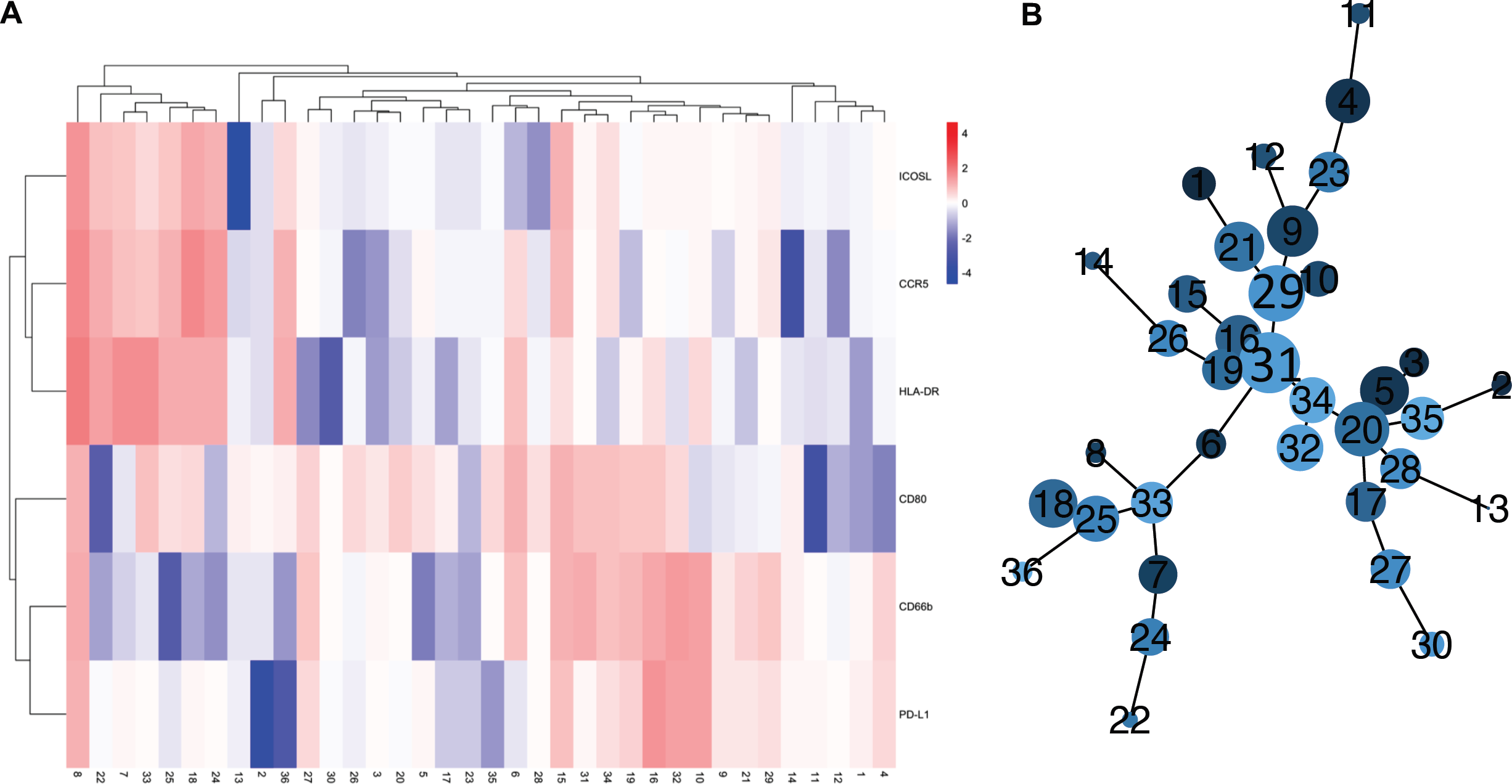
Time-course-dependent cluster analysis of PMN diversification. **A)** Hierarchical cluster heatmap depicting the expression of receptors between resting polymorphonuclear cells (rPMNs) and GM-CSF plus IFNγ- or TNF treated cells based on time-course flow cytometry data (2, 18, 24, 48, and 64 h; n=4 for each condition). The clustering was performed using CytoTree, utilizing all CD66b^+^, SSC^int^, and Zombie^-^ events. **B**) Construction of PMN diversification trajectory based on UMAP coordinates using MST and annotation according to clusters.

**Figure S5.**
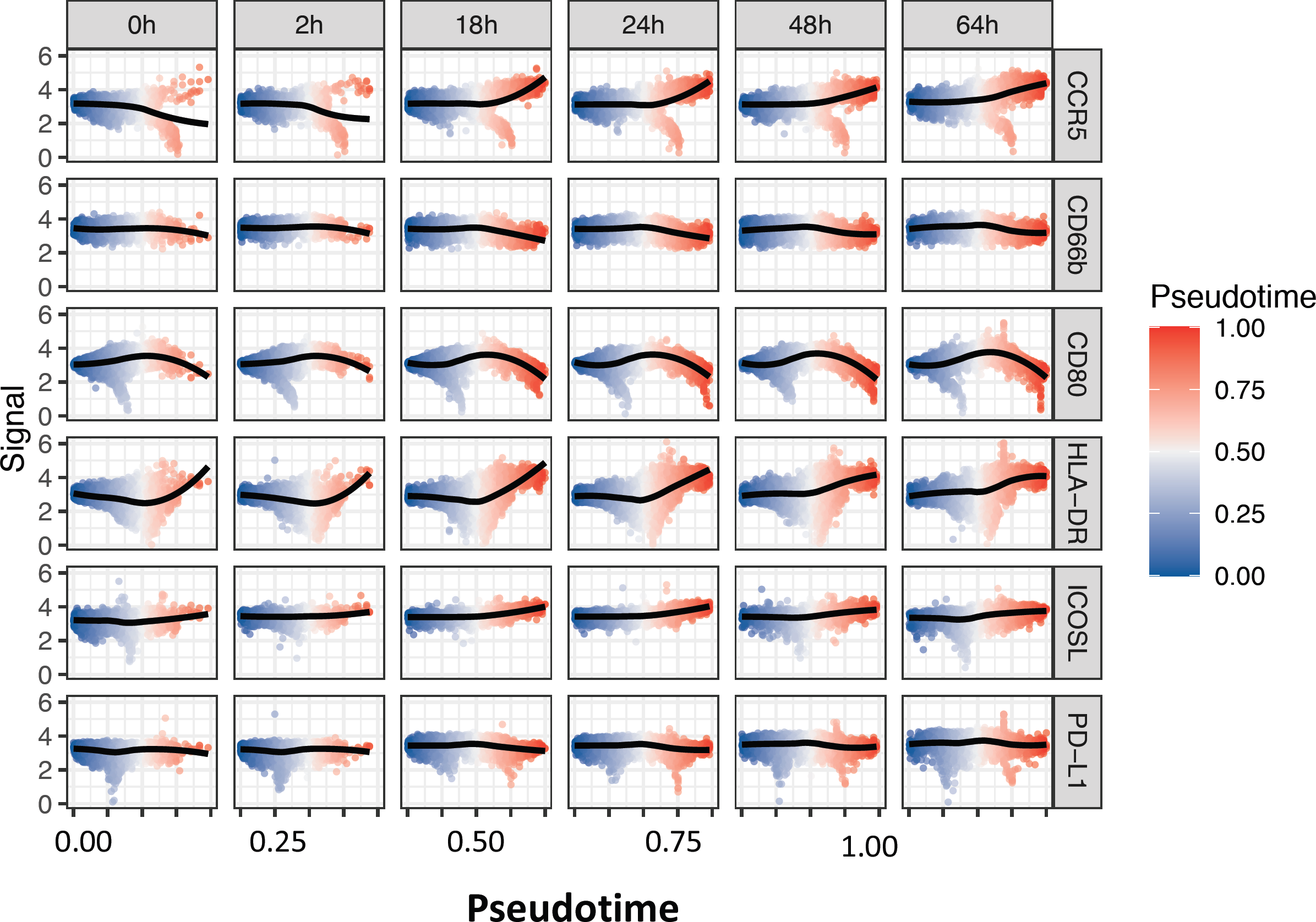
Correlation between marker expression and pseudotime in time course analysis. Pseudotime was plotted against signal strength for each marker and each time point using R/CytoTree.

**Figure S6.**
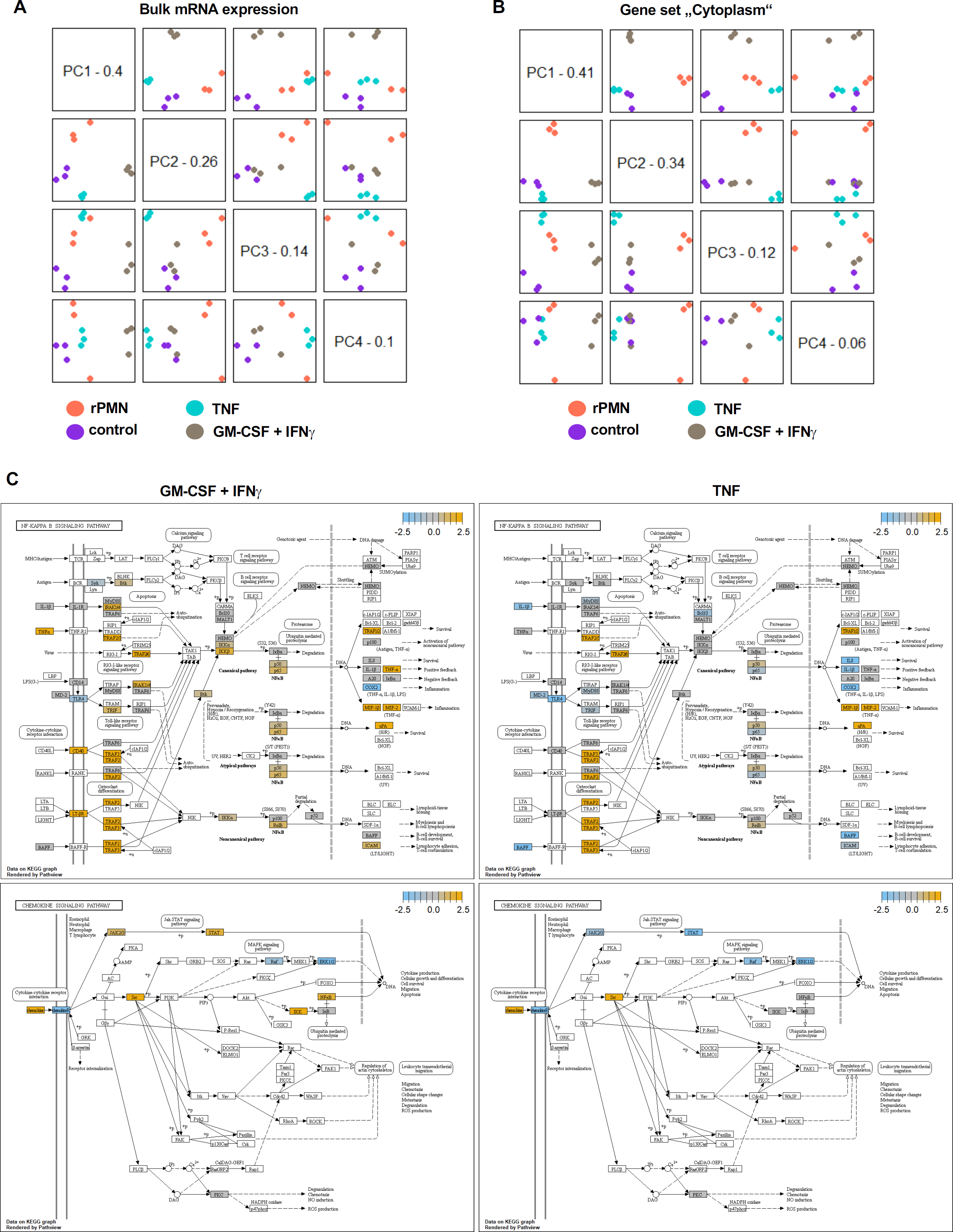
Signaling pathways in primed PMN. **A)** Principal Component Analysis (PCA) of bulk mRNA expression data of resting PMNs and primed PMNs performed using GEO: GSE194366 datasets and nCounter software. The first four principal components were plotted against each other, and colors indicate the values of each covariate. **B**) For each KEGG pathway, genes within the panel are mapped to the pathway, and differential expression information is overlaid on the protein-based KEGG pathway image. Pathway nodes shown in white have no genes in the panel that map to them. Pathway nodes in grey have corresponding genes in the panel, but no significant differential expression is observed. Nodes in blue denote downregulation relative to the selected baseline, whereas nodes in orange denote upregulation relative to the selected baseline.

**Figure S7.**
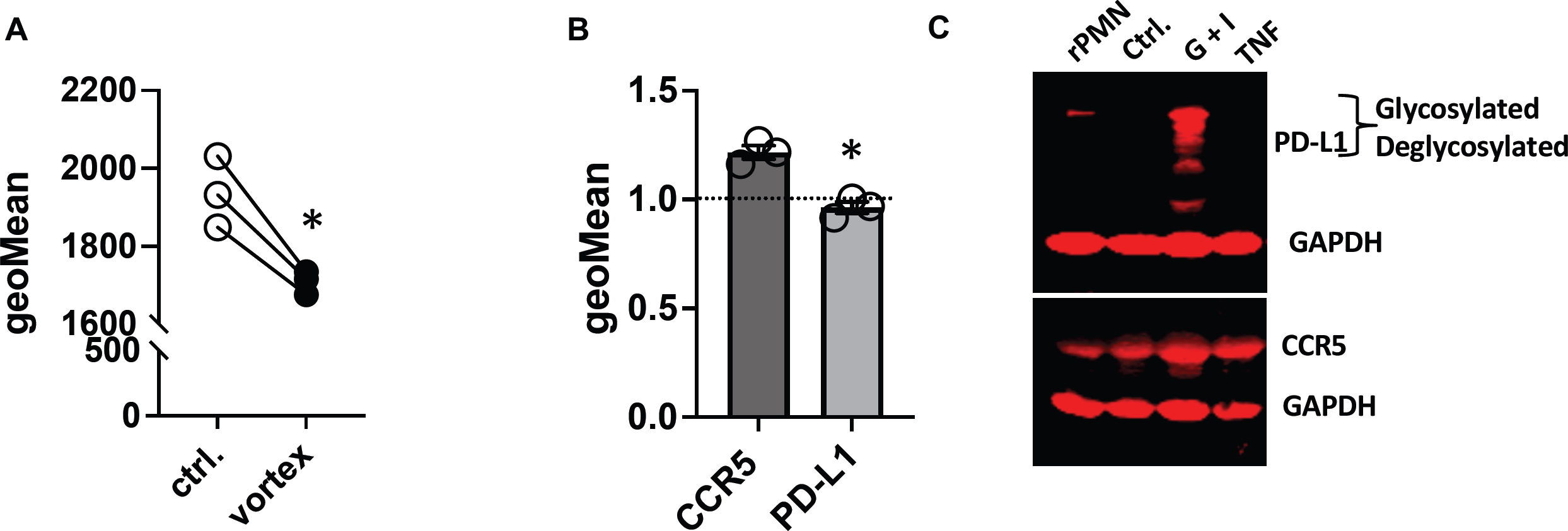
Shear force affects actin perturbation and PMN diversification. **A)** F-actin content in PMNs was measured in resting PMNs (control) or after vortex-induced mechanical stress (1 min. at 3,000 rpm) using fluorescently-labeled phalloidin (SEM, n=3, t-test, *p<0.05). **B**) CCR5 and PD-L1 expression was quantified 24 h after vortex-induced mechanical stress (1 min at 3,000 rpm). The graph shows the relative change in expression compared to untreated PMNs (SEM, n=3, t-test, *p<0.05). **C**) Whole cell lysates were prepared, and the proteins were separated using SDS-PAGE. Following the protein transfer to a membrane, detection was carried out using fluorescent antibodies as specified. GAPDH was used as a loading control on separate blots. The presented blot represents the results obtained with PMNs from three individuals.

**Figure S8.**
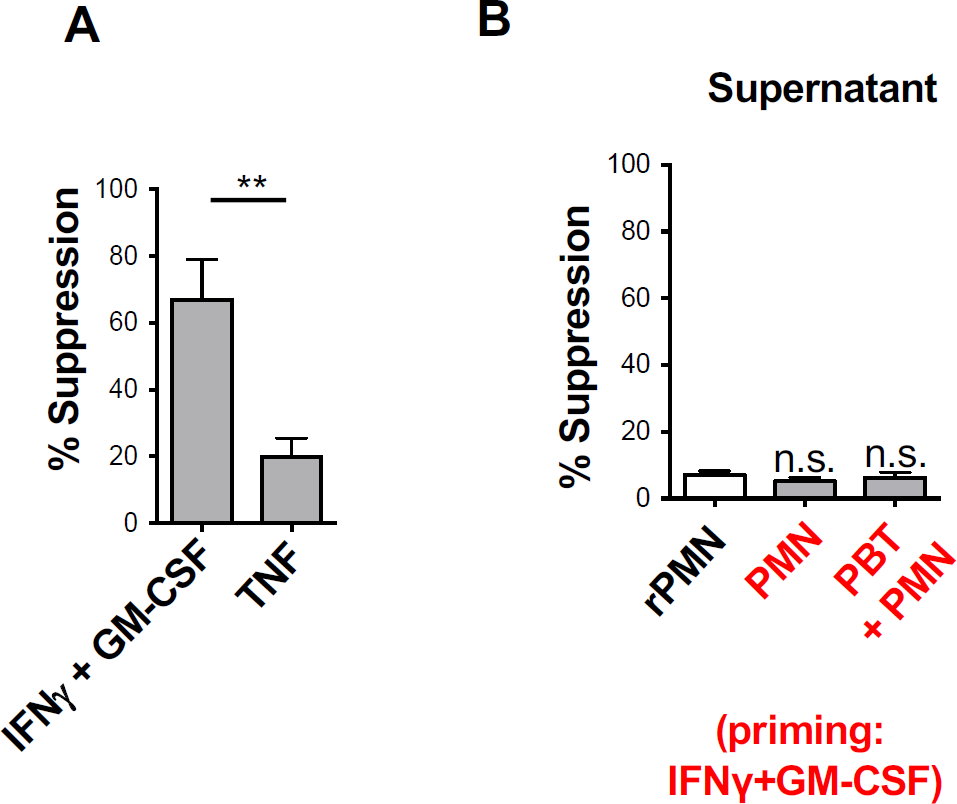
T-cell suppression assay with primed PMN or supernatants. **A**) T cells were co-stimulated and incubated with either GM-CSF + IFNγ or TNF-primed PMNs. Proliferation was measured using CFDA dilution assays and is depicted as percent suppression compared to co-stimulated cells in the absence of PMNs (n=3, **p<0.01). **B**) Inhibition of co-stimulation-dependent T cell proliferation by supernatants from cell cultures with resting PMNs, primed PMNs, or T cell/primed PMN co-cultures was measured as discussed above (SEM, n=3, n.s. = not significant).

**Table S1.**
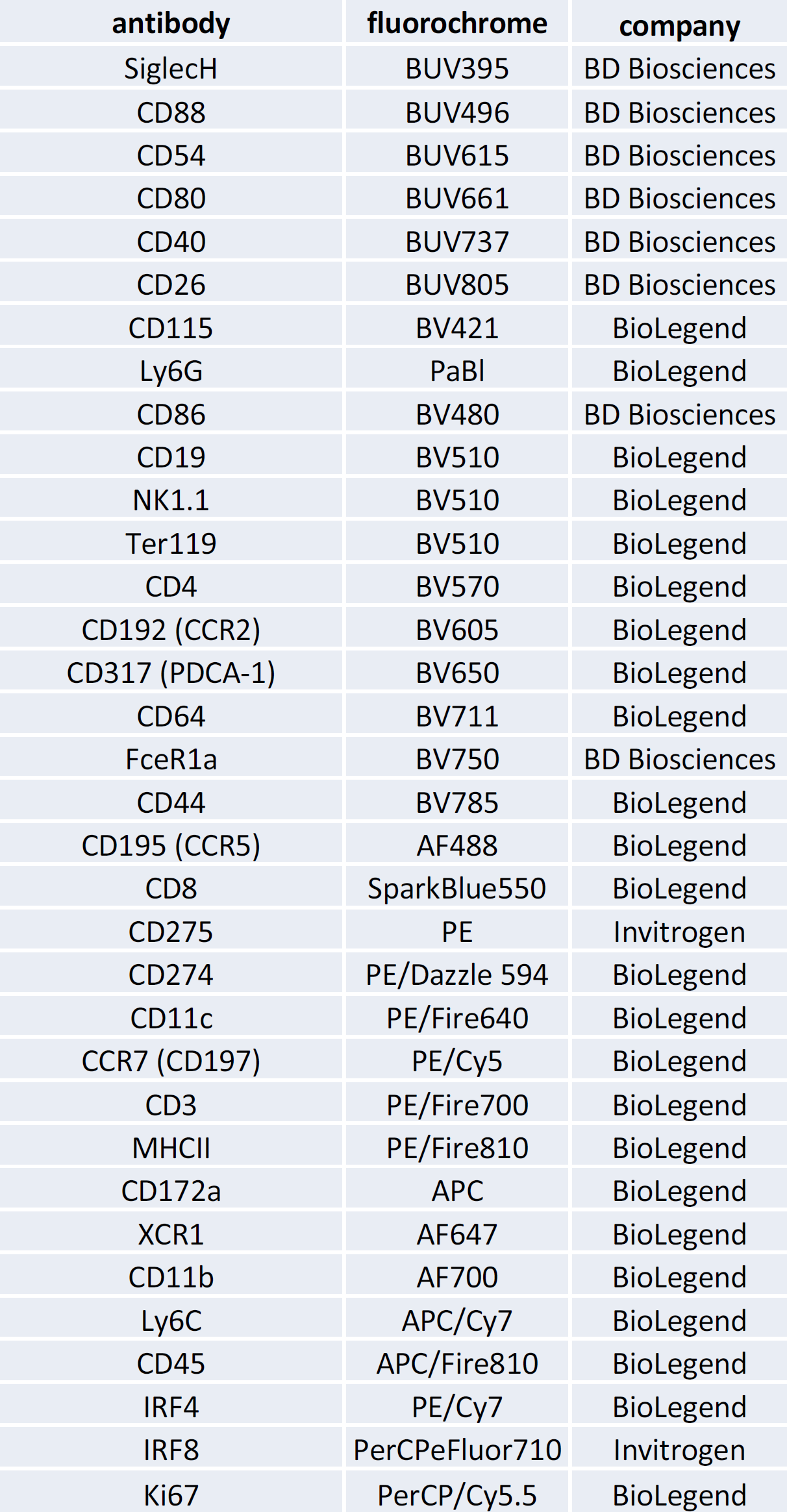
Antibody panel for flow cytometry analysis of PMNs of mice.

## Notes

There is no conflict of interest

